# Detection of interictal epileptiform discharges: A comparison of on-scalp MEG and conventional MEG measurements

**DOI:** 10.1101/834275

**Authors:** Karin Westin, Christoph Pfeiffer, Lau M. Andersen, Silvia Ruffieux, Gerald Cooray, Alexei Kalaboukhov, Dag Winkler, Martin Ingvar, Justin Schneiderman, Daniel Lundqvist

## Abstract

Magnetoencephalography (MEG) is an important part of epilepsy evaluations because of its unsurpassed ability to detect interictal epileptiform discharges (IEDs). This ability may be improved by next-generation MEG sensors, where sensors are placed directly on the scalp instead of in a fixed-size helmet, as in today’s conventional MEG systems. In order to investigate the usefulness of on-scalp MEG measurements we performed the first-ever measurements of on-scalp MEG on an epilepsy patient. The measurement was conducted as a benchmarking study, with special focus on IED detection. An on-scalp high-temperature SQUID system was utilized alongside a conventional low-temperature “in-helmet” SQUID system. EEG was co-registered during both recordings. Visual inspection of IEDs in the raw on-scalp MEG data was unfeasible why a novel machine learning-based IED-detection algorithm was developed to guide IED detection in the on-scalp MEG data. A total of 24 IEDs were identified visually from the conventional in-helmet MEG session (of these, 16 were also seen in the EEG data; eight were detected only by MEG). The on-scalp MEG data contained a total of 47 probable IEDs of which 16 IEDs were co-registered by the EEG, and 31 IEDs were on-scalp MEG-unique IEDs found by the IED detection algorithm. We present a successful benchmarking study where on-scalp MEG are compared to conventional in-helmet MEG in a temporal lobe epilepsy patient. Our results demonstrate that on-scalp MEG measurements are feasible on epilepsy patients, and indicate that on-scalp MEG might capture IEDs not seen by other non-invasive modalities.

## 1. Introduction

Magnetoencephalography (MEG) has played a role in epilepsy care for almost thirty years, and is today widely regarded as an established clinical tool (De Tiège et al., 2012; Hari et al., 2018). Several studies have demonstrated that MEG detects interictal epileptiform discharges (IEDs) with unsurpassed sensitivity, detecting them in approximately 70-80% of all epilepsy patients as compared to a 60% detection rate in EEG (Knake et al., 2006; Stefan et al., 2003). MEG also plays a role in EEG-negative epilepsy cases, and adding MEG to the clinical evaluation of such patients increase the spike detection probability with almost 20% (Colon et al., 2009; Duez et al., 2016; Pataraia et al., 2004). Furthermore, since MEG source reconstruction is less affected by skull anatomy and conductivity than EEG is, the localization of an epileptogenic zone is more accurate with MEG than what is possible with EEG (Hämäläinen et al., 1993; Jayakar et al., 2014). Also, using MEG to guide intracranial electrode placement increases the likelihood of a successful sampling of the seizure onset zone (Jung et al., 2013; Knowlton et al., 2006; Sutherling et al., 2008). Additionally, resection of findings localized with MEG increases the likelihood of post-surgery seizure freedom compared to surgery performed without MEG findings taken into account (Murakami et al., 2016; Rampp et al., 2019). For the above reasons, MEG has become a standard part of presurgical evaluation of epilepsy patients (De Tiège et al., 2017; Hari et al., 2018).

Despite these unique contributions in presurgical epilepsy evaluation, conventional MEG systems exhibit some inherent limitations, and addressing these might further enhance the utility of MEG in epilepsy research and clinical evaluations. Conventional MEG (hereafter called in-helmet MEG) sensors are cooled down to approximately 4 K (−269 °C) using liquid helium, which is why they must be housed behind thick layer of insulation (Heiden, 1991) within a fixed-size helmet. On adults, this solution results in a 20-40 mm sensor-scalp distance typically influencing distance to frontal and temporal cortices the most; the situation is even worse for children (Riaz et al., 2017). This distance has a detrimental influence on the signal-to-noise ratio of the cortical signal since the magnetic field strength weakens quickly with distance (Boto et al., 2016; Iivanainen et al., 2017). Spatial resolution depends on the sensor spacing. A smaller sensor-to-sensor distance results in a better ability to distinguish between neural sources, compared to a greater sensor-to-sensor distance (Boto et al., 2016; Riaz et al., 2017). To address both the problems of sensor-cortex distance and that of the fix in-helmet system, as well as improve both neuroscientific and clinical applicability of MEG, systems where the sensors are flexibly placed directly on the scalp are under development (Borna et al., 2017; Boto et al., 2018; Iivanainen et al., 2019; Pfeiffer et al., 2019). On-scalp MEG sensors comprise, amongst others, optically-pumped magnetometers (OPMs) (Budker and Romalis, 2007) and high-Tc SQUIDs (Zhang et al., 1993). Both of these on-scalp MEG sensor systems allow for a significant reduction of the sensor-cortex distance, as well as a rearrangement of the sensor layout geometry, thus increasing the signal-to-noise ratio and spatial resolution of the recorded neuronal activity (Boto et al., 2016; Iivanainen et al., 2017; Riaz et al., 2017; Schneiderman, 2014). Furthermore, placing the sensors evenly distributed on the scalp enables a more even sampling of brain regions. Thus, the development of on-scalp MEG sensors holds the promise of improving the quality of non-invasive MEG measurements, potentially moving these towards the quality of intracranial registrations. Potentially, on-scalp MEG sensors could enable better non-invasive characterization of focal epileptic networks, seizure development and seizure onset zone, which today is only possible using invasive intracranial recordings (Bartolomei et al., 2017; Jayakar et al., 2016, 2014; Stefan and da Silva, 2013). Improving the spatial resolution of non-invasive neurophysiological measurements would thus be of great value both for neuroscientific and clinical applications.

We present the first-ever measurement on an epilepsy patient using on-scalp MEG sensors. We aimed to evaluate the potential added value of these sensors compared to in-helmet MEG with focus on IED detection. To this end, a benchmarking protocol with acquisition of both on-scalp and in-helmet MEG and co-registration of EEG from the same patient was utilized.

## 2. Method and material

### 2.1 Ethical approval

The experiment was approved by the Swedish Ethical Review Authority (DNR: 2018/1337-31), and was performed in agreement with the Declaration of Helsinki.

### 2.2 MEG systems

#### 2.2.1 On-scalp high-T_c_-MEG system

The on-scalp high-Tc-MEG system (hereafter referred to as on-scalp MEG) consists of seven SQUID magnetometers, each with a pickup loop with dimensions 8.6 mm x 9.2 mm. The magnetometers are positioned with 12.0 mm center-to-center distance in a hexagonal array enclosed within a cryostat cooled with liquid nitrogen. The distance between the sensors and the participant’s scalp can be as small as 1 mm. A detailed description of the system is found in (Pfeiffer et al., 2019).

#### 2.2.2 In-helmet MEG system

For in-helmet MEG recordings, an Elekta Neuromag TRIUX (Elekta Oy, Helsinki, Finland) with 102 sensor chips, each with one magnetometer with a pickup loop size of 21 mm x 21 mm and two orthogonal planar gradiometers, was used.

### 2.3 Patient and experimental procedure

In order for IEDs to be feasibly detected via on-scalp recordings, they need to be focal, reliably sampled by in-helmet MEG, and frequently occurring. Furthermore, in order to compare IED properties across MEG/EEG sensors, the IED configuration should be as simple as possible, preferably distinct, solitary sharp waves or spikes. To identify potential participants that met these criteria, scalp EEG of ten adult, cognitively intact epilepsy patients who had undergone long-term video EEG as part of an epilepsy evaluation at the department of clinical neurophysiology at the Karolinska University Hospital during 2018 were screened. Six patients diagnosed with focal epilepsy were contacted; three agreed to be screened for inclusion in the benchmarking study. These three patients subsequently underwent an in-helmet MEG recording with co-registration of EEG, electrooculography (EOG), and electrocardiography (ECG). These recordings were used to identify patients with IEDs that are clearly visible on MEG. For EEG, a 10-20 montage with 21 channels was used. During measurements, patients were seated upright and instructed to try to stay awake. Data was recorded for one hour: 30 minutes with eyes closed and 30 minutes with eyes open. Two patients with prominent in-helmet MEG detected IEDs were invited to participate in further measurements involving both in-helmet and on-scalp MEG. One patient (female, 45 years old) agreed to further participation. This patient is diagnosed with left temporal lobe epilepsy and underwent epilepsy surgery in 1996, resulting in only a short period of seizure freedom.

### 2.4 Benchmarking measurements and analysis

The main measurements involved both in-helmet and on-scalp MEG measurements from the epilepsy patient and were conducted in accordance with the benchmarking protocol described by Xie et al. (Xie et al., 2017). In short, this protocol involves an initial measurement using in-helmet MEG, from which the magnetic fields related to the brain activities of interest are projected to the scalp to guide the placement of the on-scalp MEG system, currently having a small and limited scalp coverage.

#### 2.4.1 In-helmet MEG session

##### 2.4.1.1 Measurements

An initial measurement session of one hour was performed, involving in-helmet MEG with co-registered EEG, EOG, and ECG. EEG was recorded with the 21 electrodes previously mentioned, based on the 10-20 placement system. A total of 74 points of the head including the 21 EEG electrodes were digitized with a Polhemus Fastrak system. During the session, the patient was asked to rest with closed eyes, while remaining awake during measurements. Data was sampled at 5000 Hz, online low and high pass filtered at 1650 and 0.1 Hz, respectively. The EEG data was recorded together with the MEG data, using the TRIUX EEG channels.

##### 2.4.1.2 Analysis

In-helmet MEG data was initially pre-processed using MaxFilter (Elekta Neuromag) signal-space separation (Taulu and Simola, 2006) (buffer length 10 s, cut-off correlation coefficient at 0.98). The EEG signal and the maxfiltered raw in-helmet MEG data was filtered using a 1-40 Hz Butterworth bandpass filter in order to allow visual inspection of the signal. IEDs were detected via visual inspection of the in-helmet MEG and co-registered EEG data by a physician (KW) trained in IED detection both in EEG and MEG data. IEDs were averaged across events and source localization was performed using software package MNE Python (Gramfort et al., 2013). Minimum norm estimates (MNE) (Hämäläinen and Ilmoniemi, 1994) was used to localize the IED origin (Fig. 1). To this end, the patient’s clinical MRI was used to create a full head and brain segmentation using FreeSurfer (Dale et al., 1999; Fischl et al., 1999). The segmentation was used to determine skin, skull, and brain surface boundaries using the MNE-C software watershed algorithm (Gramfort et al., 2013). A source and single compartment volume conductor model based upon these were created using MNE-C. The locations of the peak positive and negative magnetic fields of the IEDs averaged across events were determined and plotted alongside the EEG electrode positions on a model of the patient’s head that included the EEG cap and 74 digitalization points (Fig. 2). These projections and points were used to guide the positioning of the on-scalp MEG sensor array at the center of both the positive and negative peak field positions on the patient’s head.

**Figure 1:**
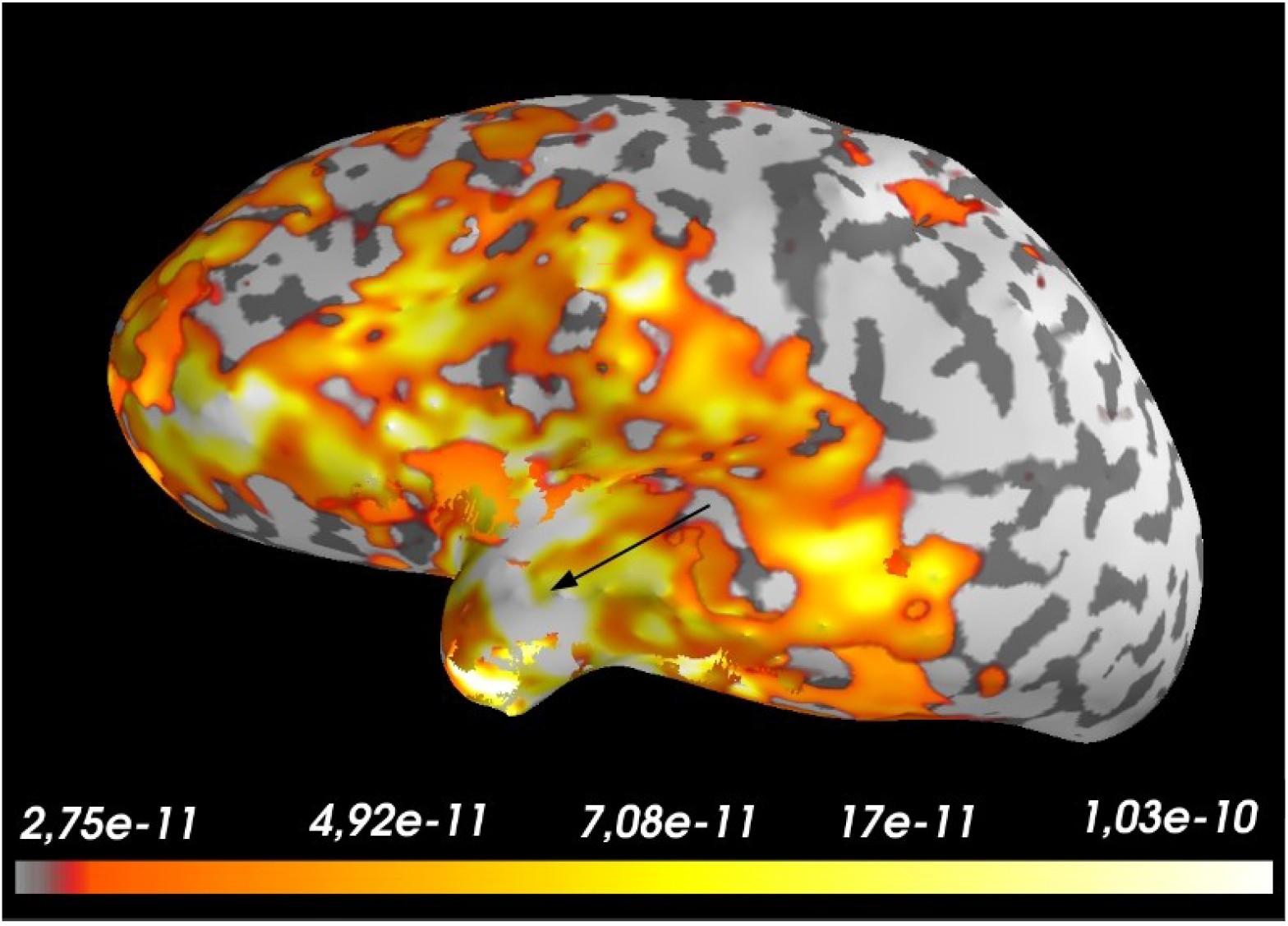
Source localization of averaged IEDs using MNE (unit: Am). Peak of IED activity marked by arrow.

**Figure 2A:**
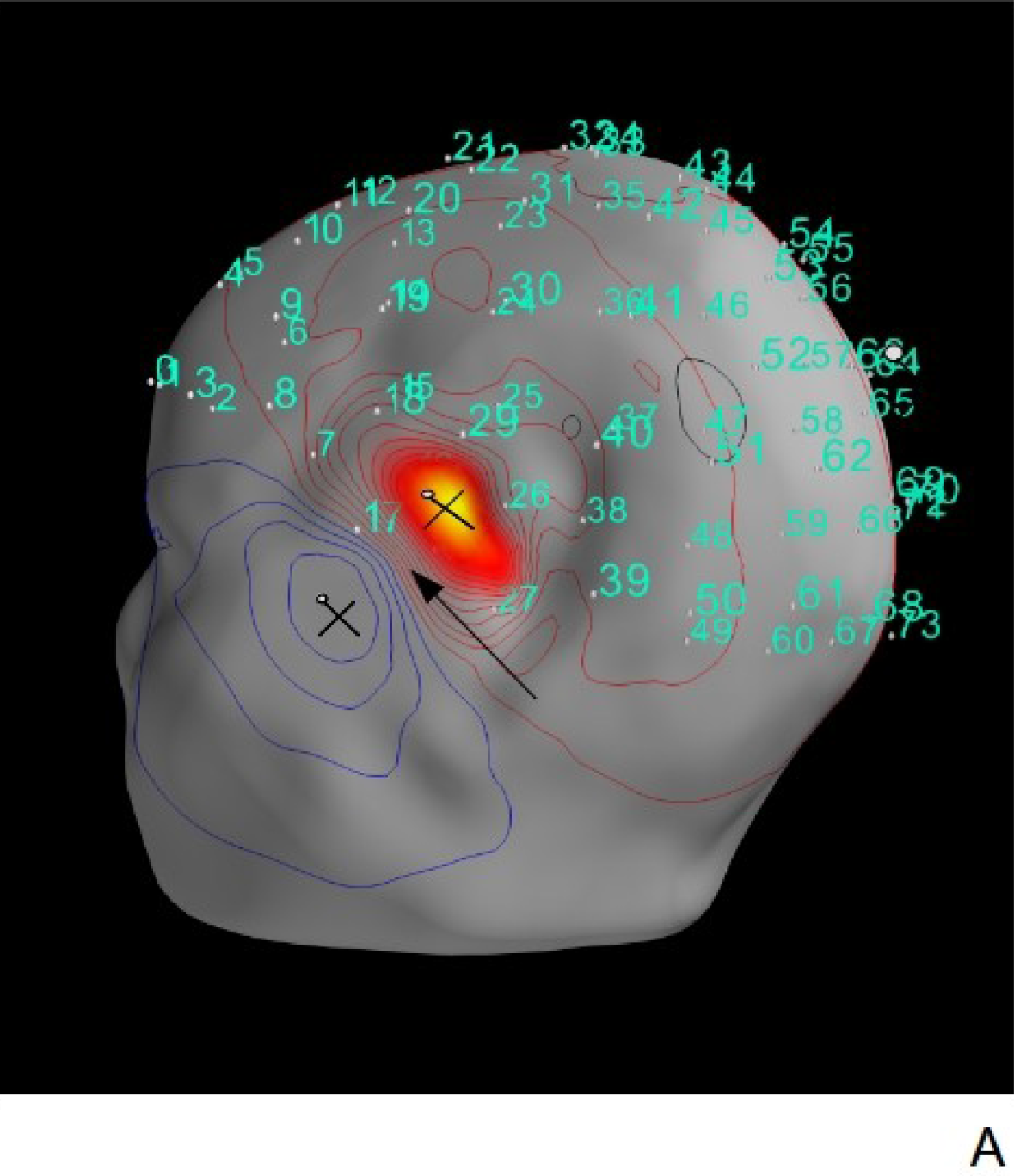
Maximum (red) and minimum (blue) peak magnetic fields of averaged IEDs and digitalized position of EEG electrodes (cyan). Peak of IED activity marked by arrow.

**Figure 2B:**
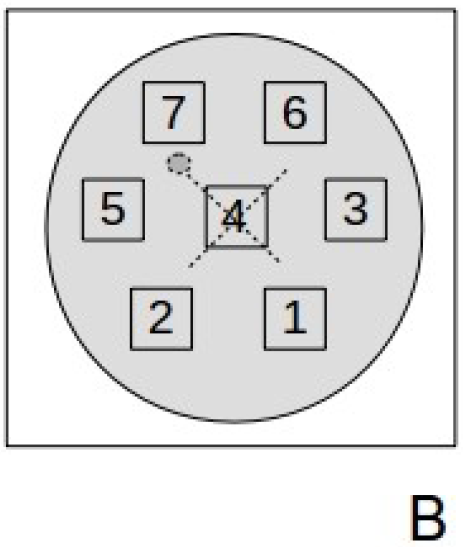
Schematic layout of on-scalp MEG sensor system. The cross indicates the positioning of the sensor at the recording sites.

#### 2.4.2 On-scalp MEG session

##### 2.4.2.1 Measurements

Two consecutive one-hour on-scalp MEG sessions were performed with the high-*T*_c_-MEG central sensor pointing at the center of each peak field position (Fig. 2). The patient was seated upright with closed eyes and asked to stay awake, similar to the in-helmet MEG measurements in Session 1. For each peak field position, co-registration of EEG was performed using the 10-20 system. During the positive peak field registration, one electrode was removed and two were slightly shifted; during the negative peak field registration, three electrodes were removed in order to make room for the on-scalp MEG system. Data from the on-scalp MEG was acquired through analog channels of the TRIUX.

On-scalp MEG data and co-registered EEG was sampled and filtered as in the in-helmet session (see *2.4.1.1, In-helmet MEG session, Measurements)*

##### 2.4.2.2 Analysis

EEG data was preprocessed as in the in-helmet MEG session. From the two on-scalp MEG recording sessions (one at the maximum field peak projected from the IEDs registered in the in-helmet MEG, one at the minimum field peak) only data from the maximum peak field recording was analyzed. Data from the minimum field peak was unfortunately rendered useless due to the removal of EEG electrodes in order to fit the cryostat, making inspection of the EEG difficult and IED detection unreliable, if not impossible. Any minimum peak on-scalp findings would hence be impossible to validate against EEG-recorded IEDs. In the data from the maximum field measurement, one high-*T*_c_ sensor was excluded due to high noise. Internal noise levels of the remaining high-*T*_c_ sensors were typically around 75 fT/Hz^1/2^ across frequencies 1-40 Hz and sensors. Visual inspection of on-scalp MEG epochs locked to IEDs in the EEG recording (hereafter referred to as EEG-positive IEDs) revealed that some of these on-scalp IEDs were sharp, transient events easily distinguishable from the background activity, while some were obscured by artifacts (Fig. 3–4). Importantly, beyond these EEG-positive on-scalp MEG IEDs, the on-scalp MEG data contained a large number of high-amplitude events visually resembling the EEG-positive IEDs (Fig 5), but without any coinciding IEDs in the co-registered EEG to validate them. Thus, visually distinguishing which of these events that might be EEG-negative, on-scalp MEG-positive IEDs, and which might be artifacts or epilepsy-related, non-IED focal activity was not possible, and an alternative approach to IED detection in this data was developed. Inspection of the dataset was performed with bandpass filtering 1-40 Hz and 5-20 Hz. Frequency bands were chosen so as to optimize visual inspection of IEDs. High-amplitude events were more distinguishable with 5-20 Hz bandpass filtering.

**Figure 3:**
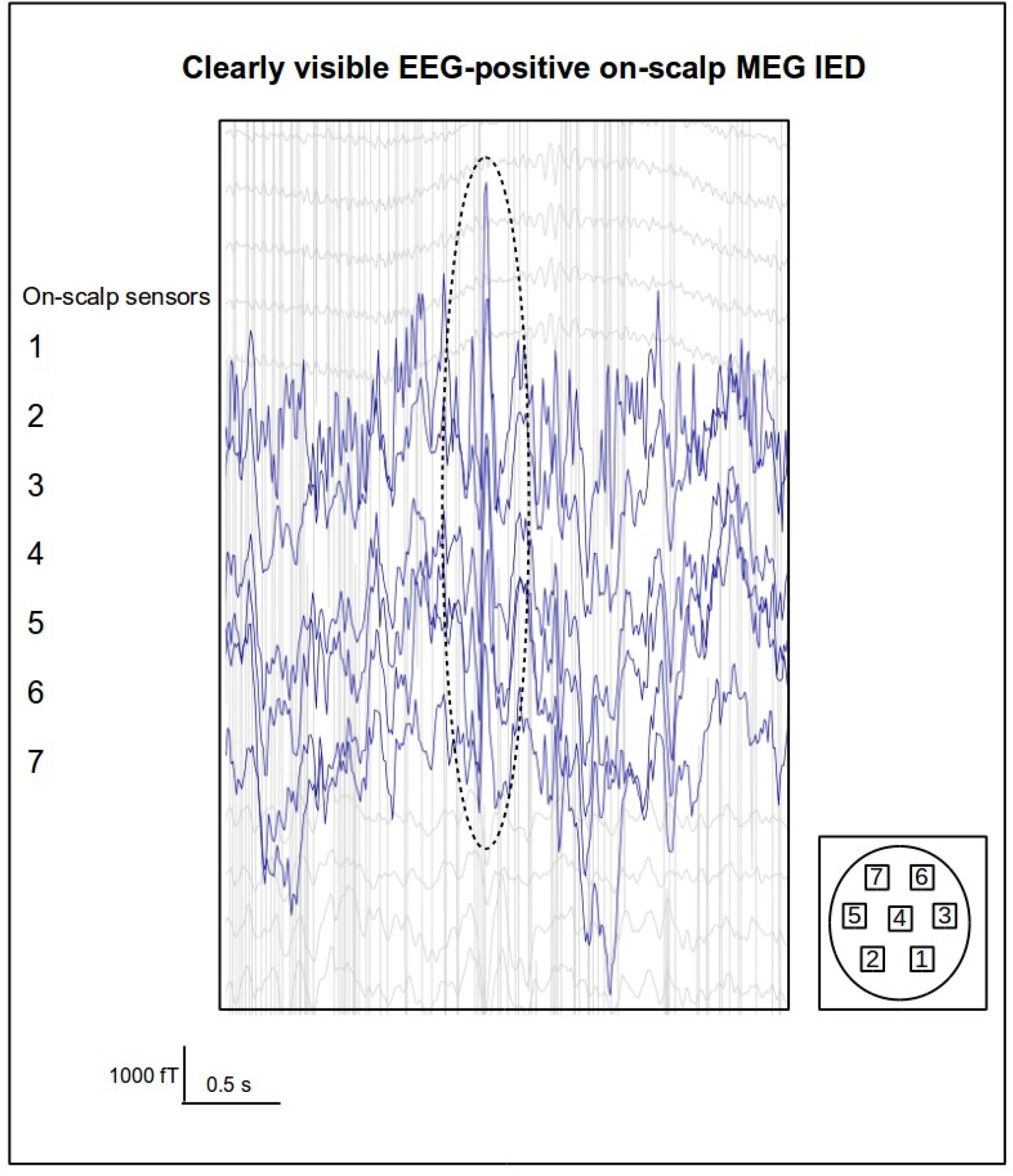
EEG-positive IED visible in raw data (bandpass filtered 1-40 Hz). On-scalp sensor numbering refer to the on-scalp MEG system layout.

**Figure 4:**
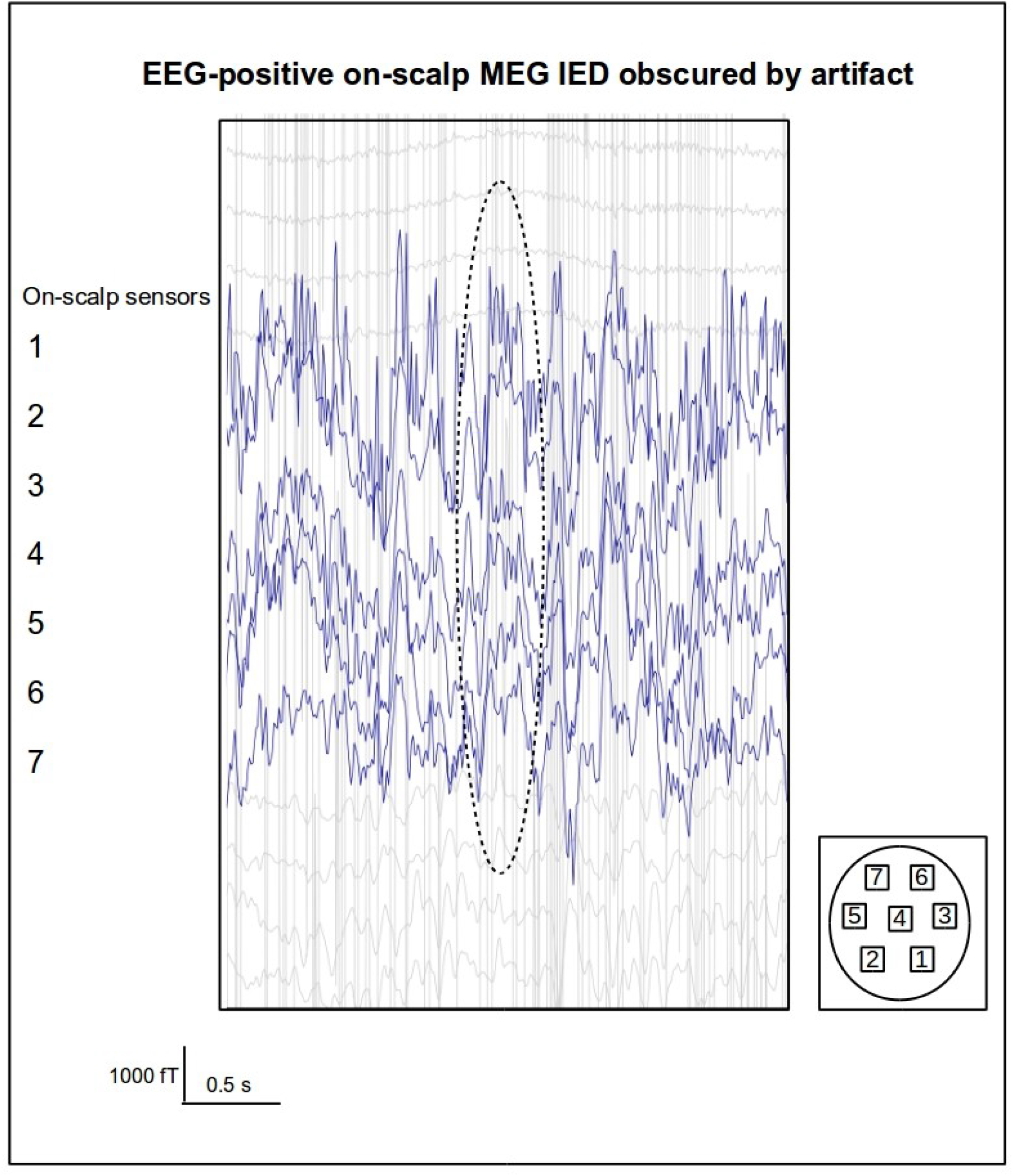
EEG-positive IED obscured by artifact (bandpass filtered 1-40 Hz). On-scalp sensor numbering refer to the on-scalp MEG system layout.

**Figure 5:**
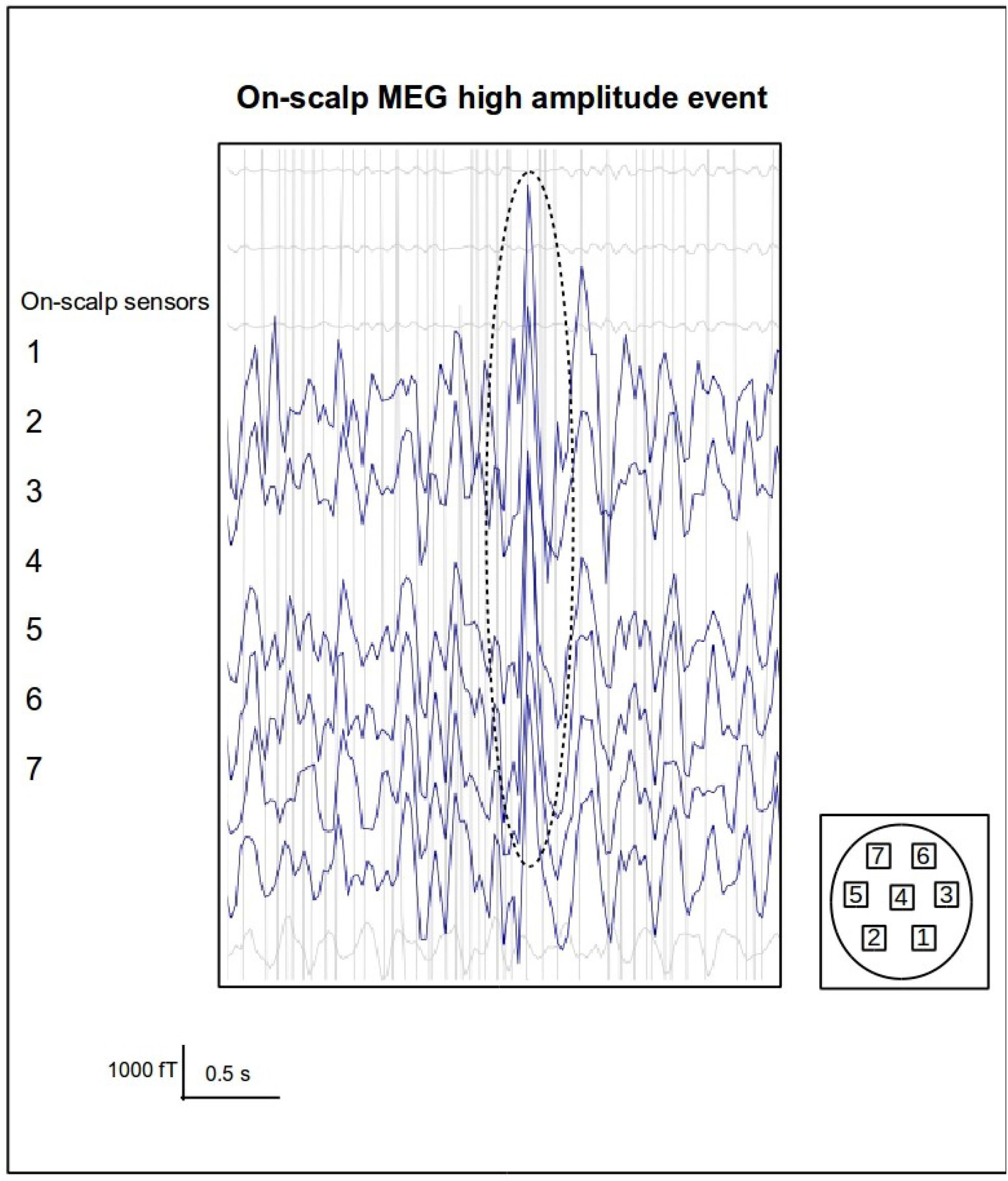
High-amplitude on-scalp MEG event (bandpass filtered 5-20 Hz). On-scalp sensor numbering refer to the on-scalp MEG system layout.

### 2.5 Spike detection

#### 2.5.1 In-helmet MEG session data

IED detection was performed as described in 2.4.1.2 *In-helmet MEG session, Analysis*.

#### 2.5.2 On-scalp MEG session data

In order to reveal whether the on-scalp MEG raw data contained any EEG-negative IEDs, a detection algorithm based upon inherent data characteristics of the on-scalp MEG data was needed. However, it is not initially given what data parameters should be used to distinguish on-scalp IED events. Definitions of interictal activity are largely arbitrary descriptions of scalp EEG-IED morphology, which varies greatly between patients (Kane et al., 2017). In order to capture their appearance, IED-detection algorithms typically depend on feature extraction from a large number of IEDs followed by classification, which can be performed using machine learning, template matching, or independent component analysis, amongst others (Wilson and Emerson, 2002). Here, we aimed to employ a similar approach combined with anomaly detection. Due to the expected difference in on-scalp and in-helmet MEG data characteristics, in-helmet MEG IEDs could not be used as templates. The one existing on-scalp recording (our current recording) therefore had to be used both for parameter extraction and spike detection validation. In order to minimize overfitting, a genetic algorithm (GA) was used to create artificial data parameter vectors resembling the corresponding real on-scalp IED data parameters.

First, on-scalp MEG IEDs time locked to IEDs found by visual inspection of the EEG-recording were located. The parameters of Table 1 were extracted from these EEG positive on-scalp MEG IEDs creating *IED feature vectors*. The genetic algorithm was used to generate *artificial IED feature vectors* resembling these. *Non-IED feature vectors* were obtained by extraction Table 1 parameters from IED-free raw data. *Artificial IED feature vectors* and *non-IED feature vectors* were used to train a support vector machine (SVM). Secondly, the SVM was evaluated on the EEG-positive on-scalp MEG IEDs, calling correctly classified ones “true positives”, and incorrectly classified ones “false negatives”. Third, classification was performed on the remaining on-scalp MEG raw data set. Positive peaks of each wave constituted the center of an epoch from which a feature vector was extracted, and classification was performed upon these vectors. Thus, on-scalp MEG events with similar statistical properties as the EEG-positive on-scalp MEG IEDs will be found (and called *potential IEDs*).

**Table 1:**
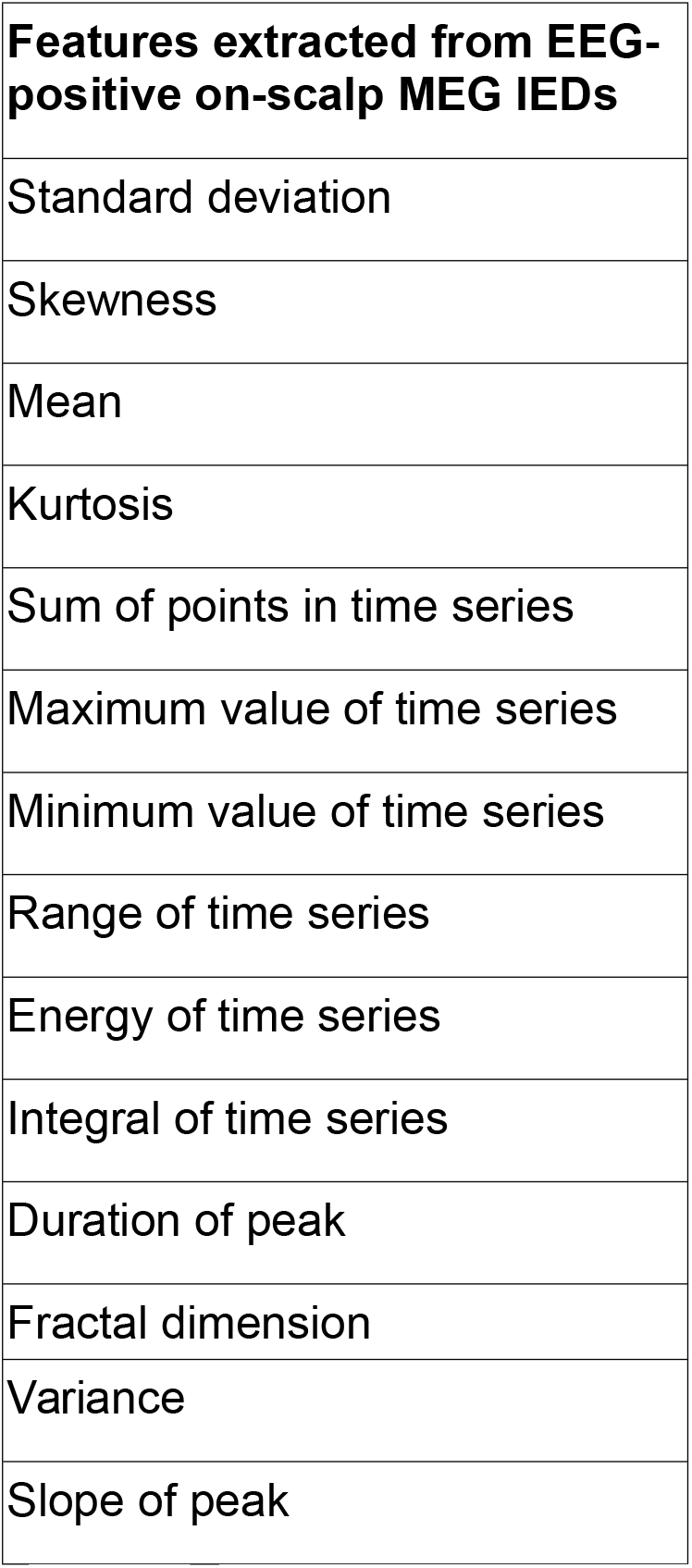
Features extracted from EEG-positive IEDs used create artificial IED feature vectors (for details, see Supplementary)

Since it is central to the IED definition that such an activity should stand out against the background activity, an IED can be considered a time series anomaly (Chandola et al., 2009; Kane et al., 2017). Only potential IEDs constituting discordant events should be kept. To this end, changes in the extracted parameters induced by the potential IEDs were quantified and only events exhibiting an equal or larger change than the smallest change exhibited by the EEG-positive on-scalp IEDs were kept. These were labeled *likely IEDs*. (For details, see Supplementary).

## 3. Results

### 3.1 In-helmet MEG session

#### 3.1.1 EEG data

From the EEG data co-registered with the in-helmet MEG recording, a total of 16 IEDs were identified via visual inspection.

#### 3.1.2 In-helmet MEG data

Visual inspection of the in-helmet MEG data revealed 24 IEDs. While 16 of these coincided with the EEG IEDs, the remaining eight in-helmet MEG IEDs were not visible in the EEG data. MNE source localization of averaged IEDs placed the epileptic focus of the MEG IEDs in the left temporal lobe (cf. Fig. 1). Amplitude of averaged IEDs was 2000 fT MNE source localization and an average of the 16 EEG-positive IEDs (i.e. the IEDs detected both in MEG and EEG data) are found in Figure 6. A corresponding MNE source localization and averaged data for the EEG-negative IEDs (i.e. the IEDs detected only in the in-helmet MEG data) are found in Figure 7.

**Figure 6:**
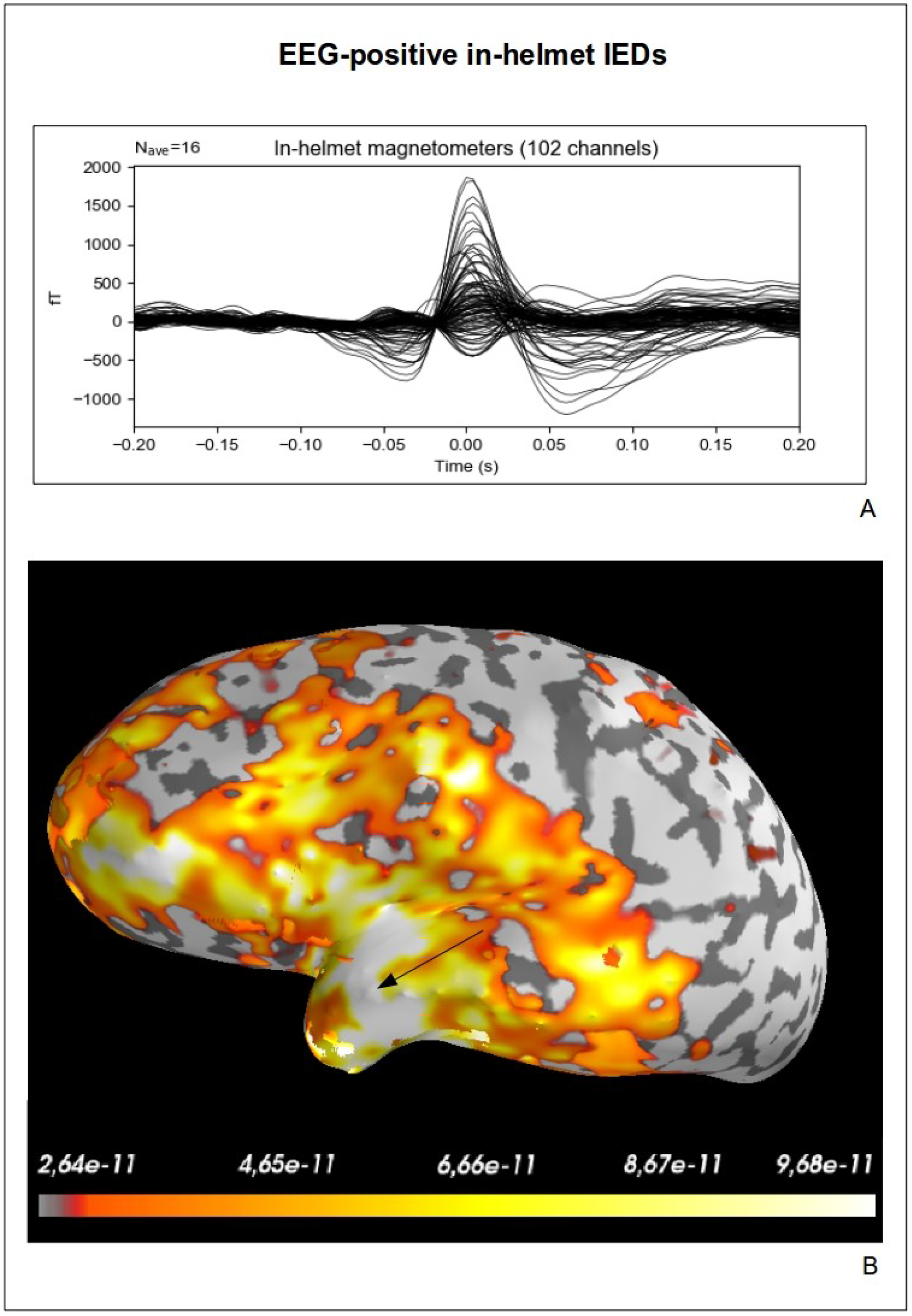
Average (A) and source localization (B) of EEG-positive IEDs found in in-helmet MEG. Source localization performed using MNE (unit: Am).

**Figure 7:**
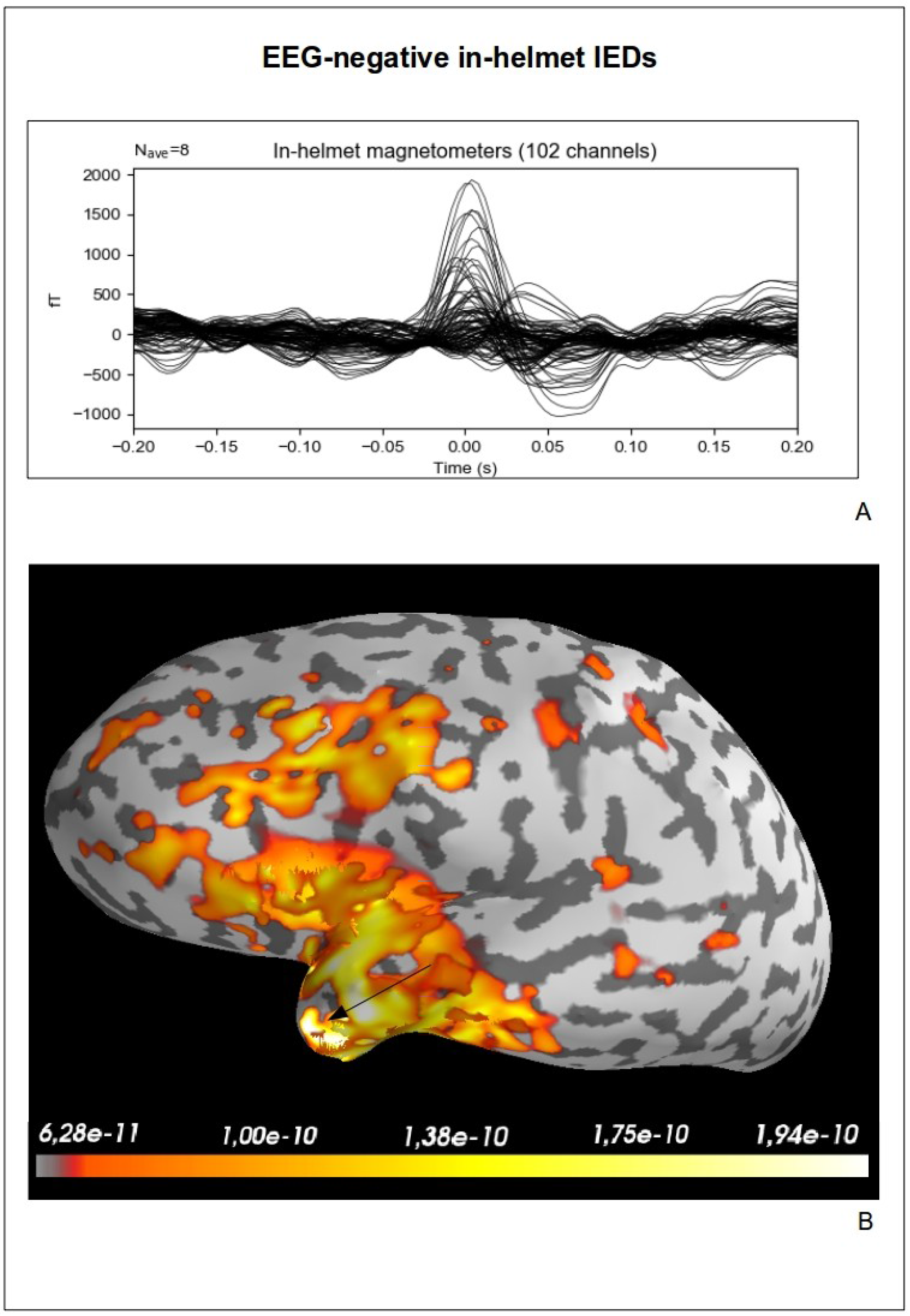
Average (A) and source localization (B) of EEG-negative IEDs found in in-helmet MEG. Source localization performed using MNE (unit: Am).

### 3.2 On-scalp MEG session

#### 3.2.1 EEG data

From the EEG data co-registered with the on-scalp MEG recording, a total of 16 IED events were detected in left temporal lobe channels, similarly to in the in-helmet MEG recording (see Fig. 8 for raw trace examples of IEDs).

**Figure 8:**
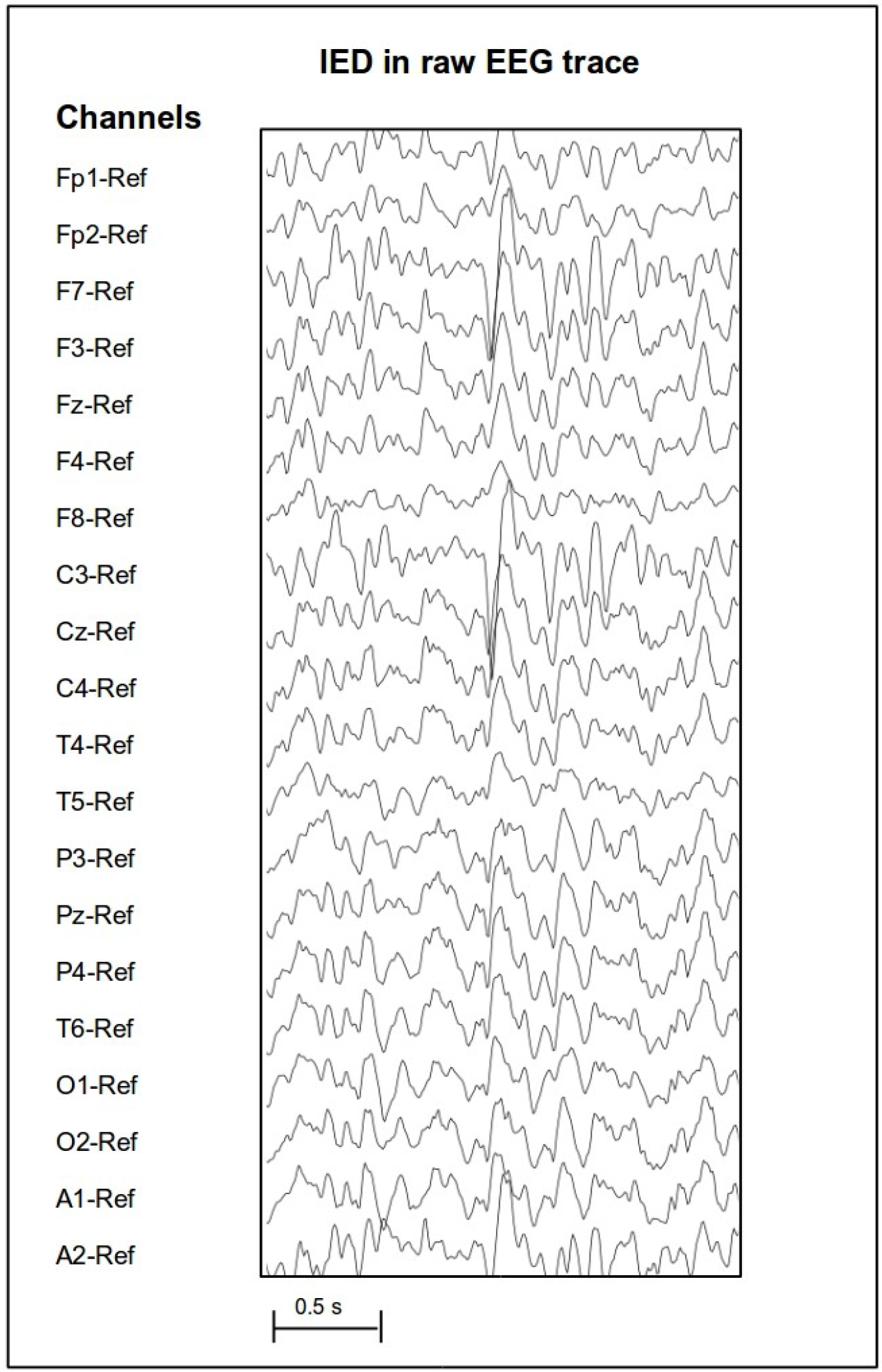
Raw trace of IED from EEG (reference montage) co-registered with on-scalp MEG.

### 3.3 Spike detection algorithm

First, the combined genetic algorithm-support vector machine (GA-SVM) was evaluated on the 16 EEG-identified IEDs events in the on-scalp MEG data (see Fig. 9 for average, see Fig. 3–4 for example of raw IEDs). Amplitude of averaged such EEG-identified on-scalp IEDs was 4000 fT. Of these, 11 events were correctly classified. Inspection of the 5 false negative epochs revealed that these on-scalp MEG events contained artifacts obscuring the IED. See Figures 10 and 11 for an average of the true positive and false negative EEG-positive on-scalp MEG IEDs; see Figure 4 for an example of a raw false negative event. Second, the GA-SVM was used to detect potential IEDs in the raw on-scalp MEG data. A total of 4623 epochs were extracted, as described in Supplementary, from the part of the on-scalp MEG recording on which classification was performed. Out of these, 416 events were classified as *potential* IEDs. An average of these are found in Figure 12.

**Figure 9:**
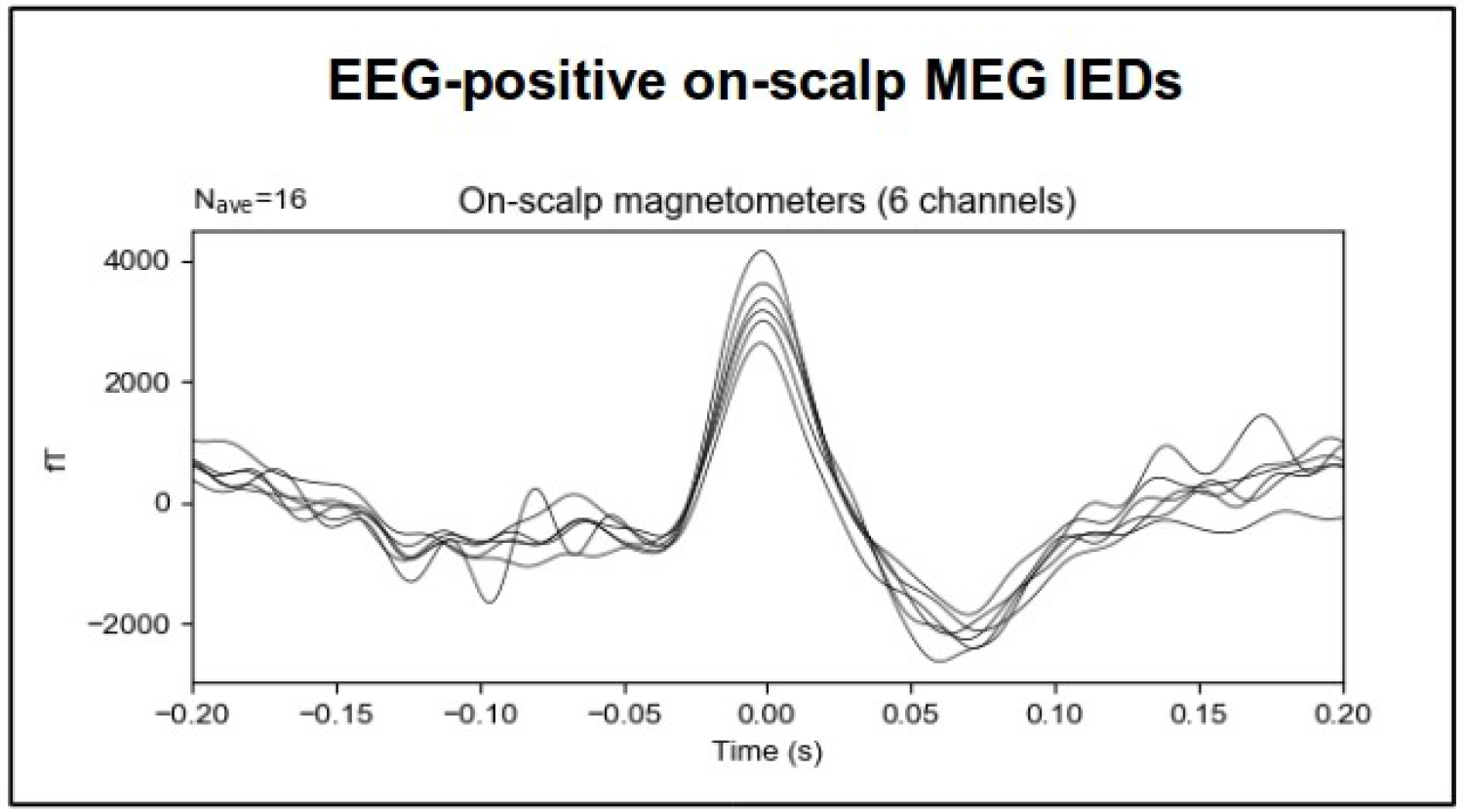
Average of all EEG-positive on-scalp MEG IEDs

**Figure 10:**
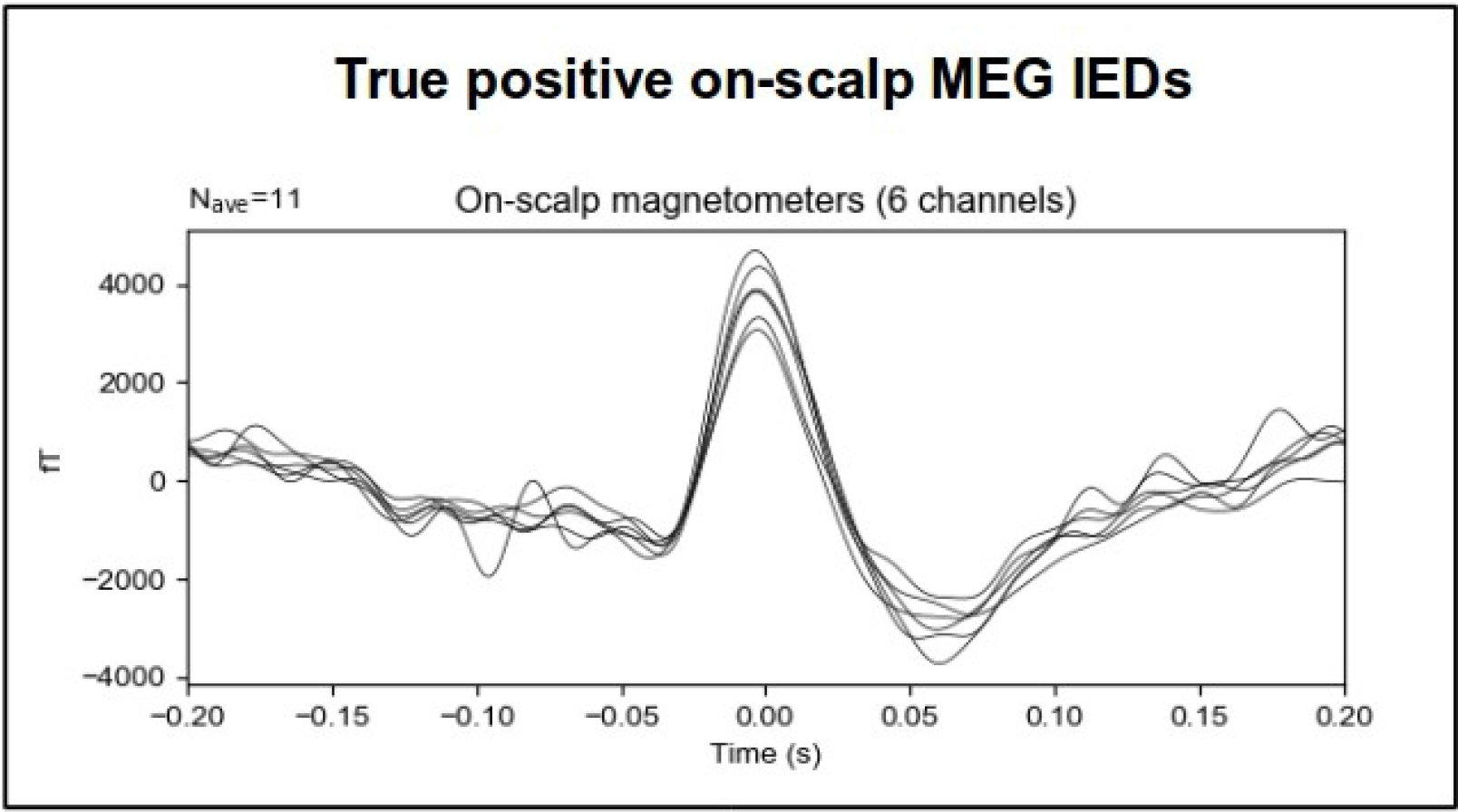
EEG-positive IEDs correctly classified as such (true positives)

**Figure 11:**
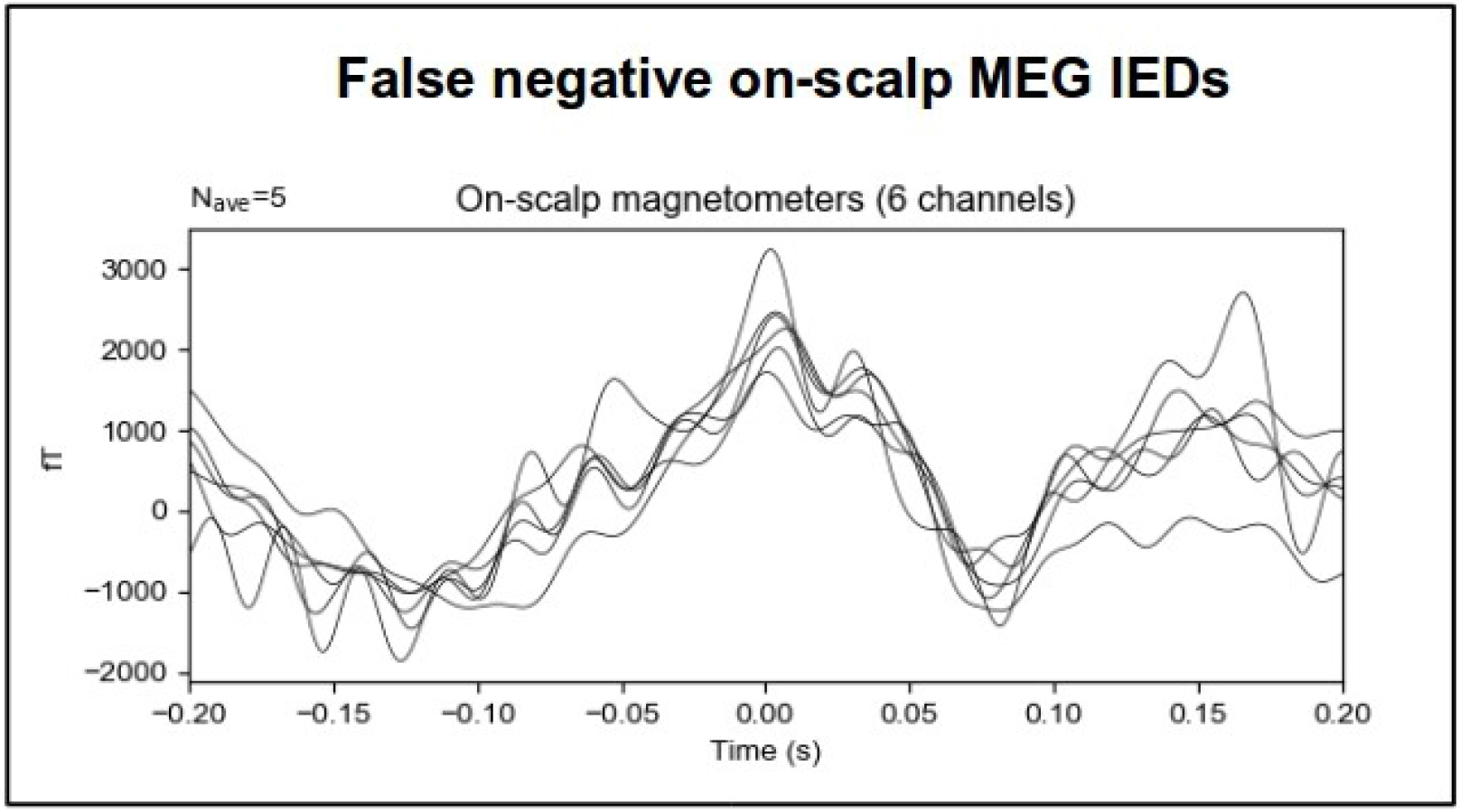
EEG-positive IEDs incorrectly classified as such (false negatives)

**Figure 12:**
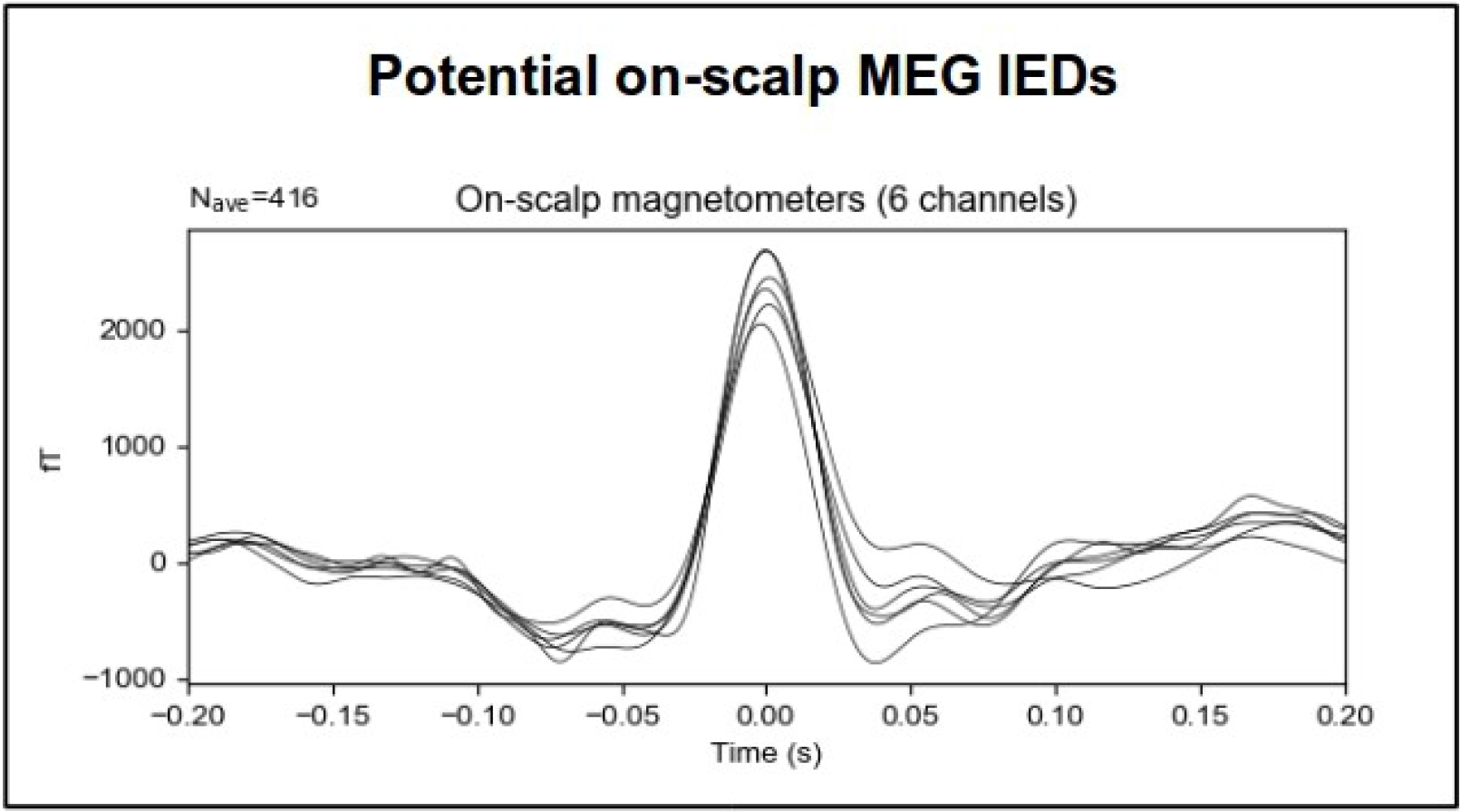
Average of potential IEDs found by the GA-SVM

Third, the potential IEDs constituting anomalies (see Supplementary for details) were kept and considered as *likely IEDs*. The on-scalp MEG recording contained 31 such likely IEDs not seen by the co-registered EEG (see Fig. 13 for average of these, and Figs. 14A,B for examples of such events in raw data). Amplitude of the averaged likely IEDs was 3000 fT.

**Figure 13:**
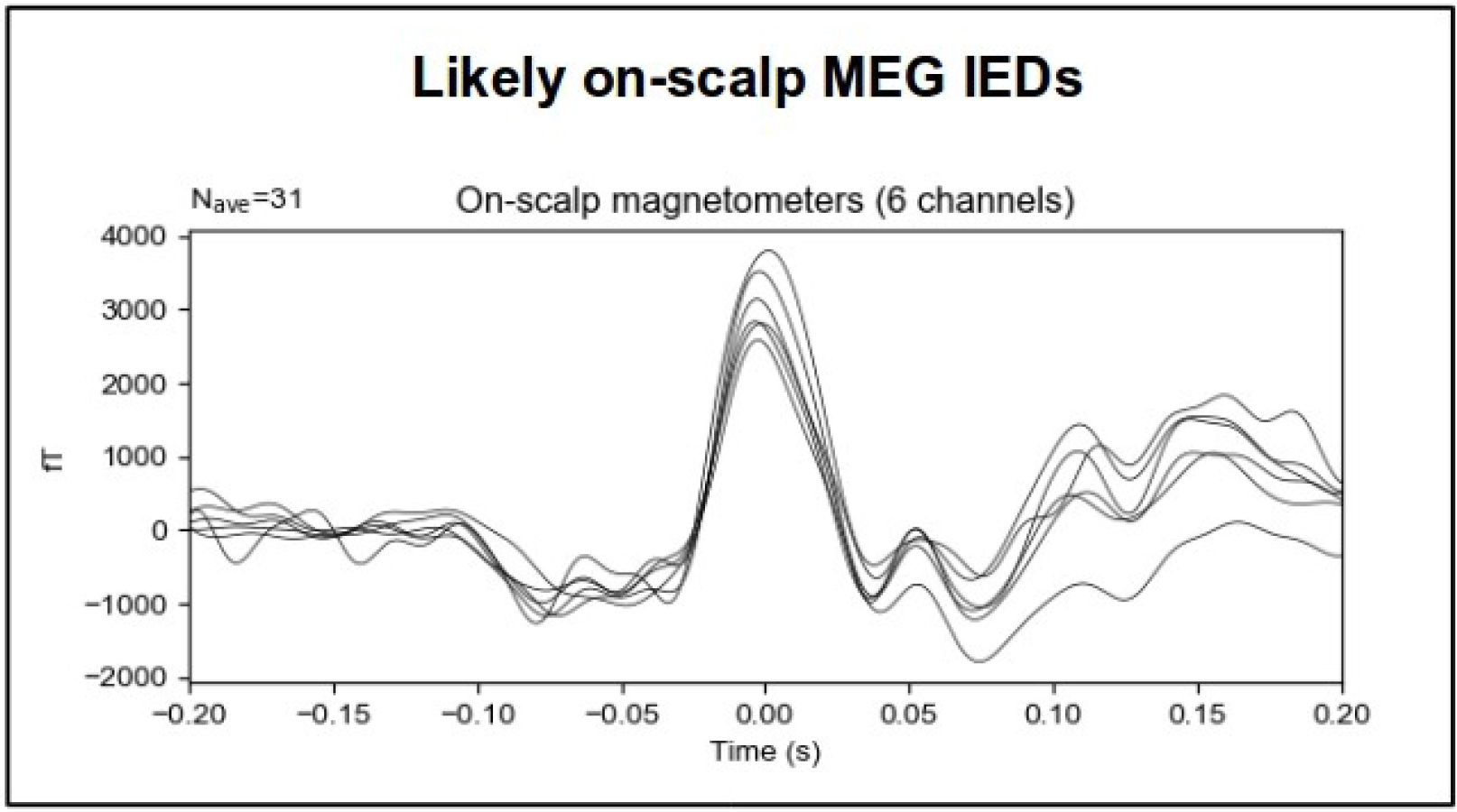
Average of likely IEDs found by combining the GA-SVM and the anomaly detector

**Figure 14A-H:**
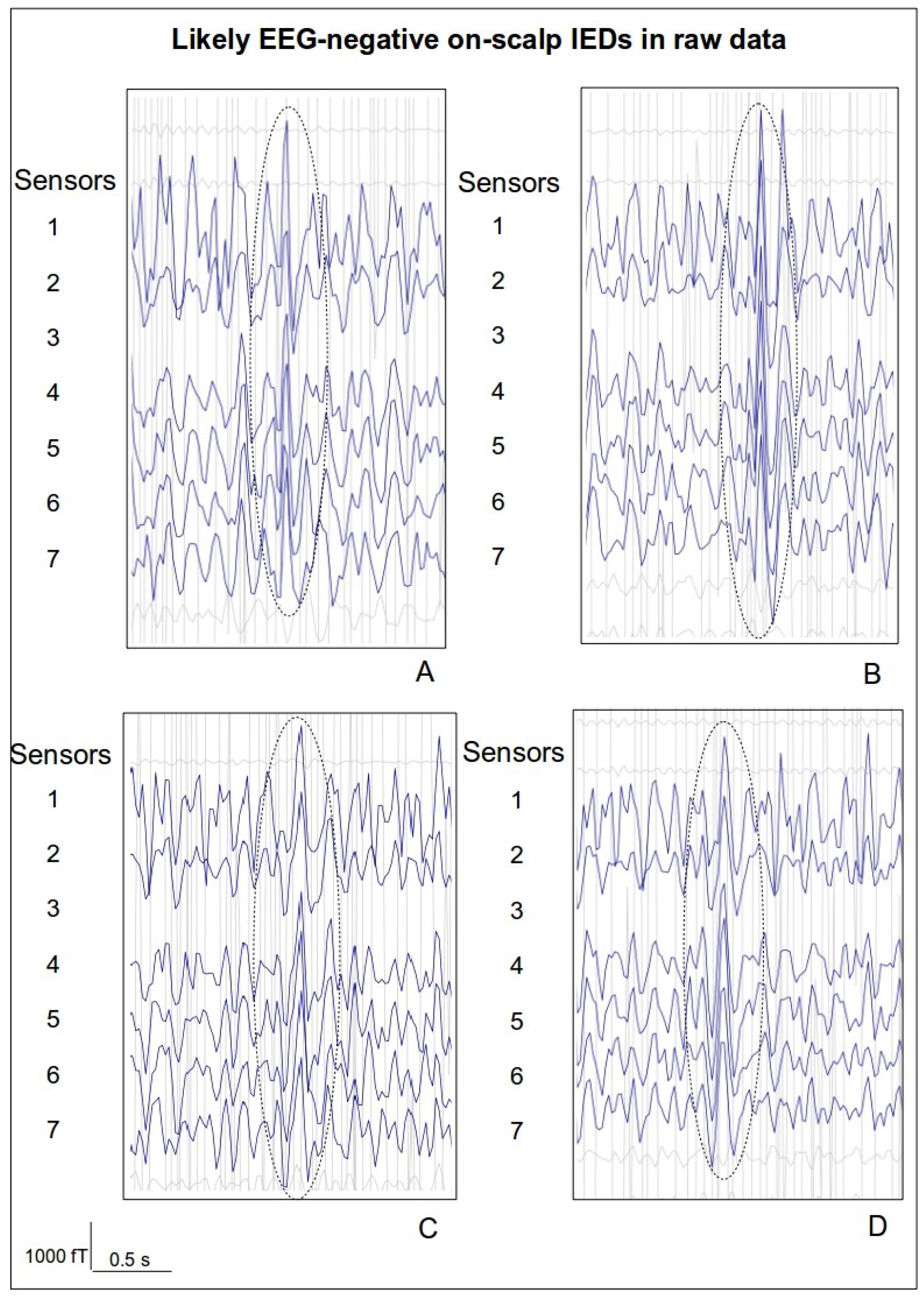
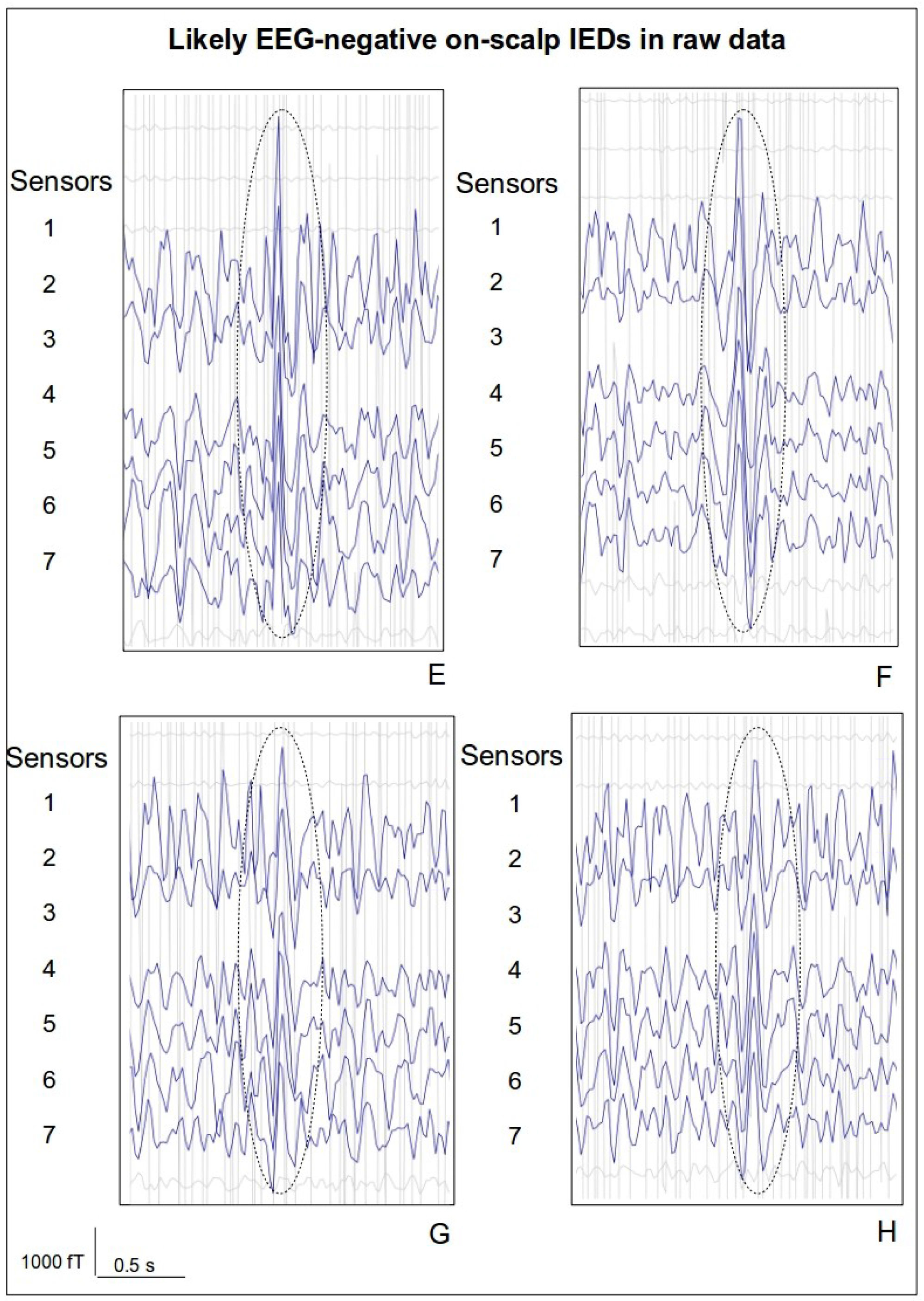
Examples of likely IED in raw on-scalp MEG data (bandpass filtered 5-20 Hz).

## 4. Discussion

We present the first-ever on-scalp MEG epilepsy study with the aim to investigate whether the sensor technology could improve non-invasive IED detection. Both on-scalp and in-helmet MEG, with co-registered EEG, was recorded from the same temporal lobe epilepsy patient. A novel on-scalp MEG IED-detection algorithm was also developed to help discern IEDs from the on-scalp MEG background activity. Below, the following aspects are discussed separately: (4.1) the feasibility of benchmarking recordings on epilepsy patients, (4.2) registration of IEDs, and (4.3) the usefulness of a detection algorithm for on-scalp MEG data.

### 4.1 Benchmarking protocol/on-scalp measurement

Data presented in this study rely on an initial careful screening of suitable epilepsy patients, and the development of a reliable benchmarking protocol. From the perspective of a study protocol, we deem on-scalp MEG recordings of epileptogenic foci activity feasible, but for now limited to patients capable of adhering to the benchmarking protocol. The temporal lobe epilepsy patient included herein exhibits relatively frequent IEDs, which enables source localization from the in-helmet MEG recording, and thus accurate placement of the on-scalp MEG system for sampling of the maximal field generated by the IEDs. However, epileptogenic foci that are difficult to localize by EEG or in-helmet MEG would hinder such optimal positioning of the on-scalp MEG system and provide poor benchmarking data, at least in studies using an on-scalp MEG array with limited coverage as we do here. A limited-coverage on-scalp MEG system is thus unlikely to be suited for measurements on patients with inconclusive non-invasive recordings, thus requiring intracranial measurements for localization (Gonzalez-Martinez et al., 2014; Jayakar et al., 2014).

### 4.2 Registration of IEDs

EEG data was co-registered with MEG in both the in-helmet and on-scalp MEG recordings. From each EEG data set, we could successfully detect 16 IEDs using visual inspection, indicating that the occurrence rate of IEDs in the patient were the same during both recordings. From the in-helmet MEG data, we could independently detect the same 16 IEDs found in the EEG data, plus an additional 8 MEG-positive IEDs. This is in line with the literature on IEDs in MEG and EEG, where MEG typically demonstrates a higher sensitivity to IEDs (Knake et al., 2006; Stefan et al., 2003).

While the IEDs in on-scalp MEG data could not be readily discriminated from other high-amplitude activities using visual inspection alone, we could guide the visual identification of the IEDs using the co-registered EEG data to validate on-scalp MEG IEDs. Leaning on EEG, we could thus identify 16 IEDs also in the on-scalp MEG data. When averaged, these on-scalp MEG IEDs revealed a prominent peak followed by hyperpolarization (Fig. 9), just like those extracted from conventional MEG measurements (Figs. 6–7). They showed typical characteristics of IEDs once we knew where they were, but were too difficult to reliably discern from other events in the data using vision alone. To explore the on-scalp MEG data for additional IEDs, we therefore used an IED detection algorithm that focuses on the abstract statistical features of IEDs, rather than their characteristic visual appearance. Using this approach, we could detect EEG-positive IEDs not obscured by artifacts (cf Fig. 4, Fig. 11 for IEDs with artifact, cf Fig. 5, Fig. 10 for clearly visible IEDs), plus an additional 31 additional IEDs uniquely registered in the on-scalp MEG data (Figs. 13–14).

Inspection of in-helmet MEG IEDs and EEG-positive on-scalp MEG IEDs (cf Fig. 6–7, Fig. 9) reveal that the field magnitude of these on-scalp MEG IEDs were roughly two times larger than the amplitude of in-helmet MEG IEDs (4000 fT and 2000 fT, respectively). This increase is in accordance with modeling predictions of the field strength acquired through a one-channel system employing the same type of on-scalp sensor used here (Xie et al., 2017, 2015). The amplitude of the 31 additional on-scalp MEG IEDs on the other hand exhibit a lower amplitude (3000 fT, Fig. 13) than do the EEG-positive ones (Fig. 9). These amplitude differences may explain why on-scalp MEG can detect IEDs that are not identified by EEG. Tao et al. have reported that the majority of IEDs visible on scalp EEG arise from hypersynchronization of at least 10 cm^2^ cortex, and no IED originating from cortical patches smaller than 6 cm^2^ can be detected with scalp EEG. However, the majority of IEDs recorded intracranially arise from smaller areas and remain undetected by scalp EEG (Tao et al., 2005). Indeed, the region capable of generating IEDs, the irritative zone (Jehi, 2018; Rosenow and Luders, 2001), is organized in subregions which might independently generate epileptic activity (Janca et al., 2018; Keller et al., 2010; Sabolek et al., 2012; Wilke et al., 2011; Zhang et al., 2017). It is thus possible that the additional IEDs detected in on-scalp MEG arise from such functional subunits of an epileptic network. Today, characterization of functional connectivity within a small region is impossible using EEG or in-helmet-MEG (Schoffelen and Gross, 2009). However, it is possible that the improved source separation and neural signal amplitude of the on-scalp MEG measurement (Boto et al., 2016; Riaz et al., 2017) would allow not only identification of such subunits, but also characterization of network dynamics. These are of course issues that need to be further explored in future on-scalp MEG measurements on epilepsy patients.

### 4.3 Using algorithm-based IED detection

Since conventional visual IED identification was unfeasible in the raw on-scalp MEG data, a SVM based IED detection was employed instead. To compensate for having only a single on-scalp MEG data set, a genetic algorithm was utilized to generate a synthetic training data set for classification based upon statistical features of the EEG-locked on-scalp MEG IEDs. The results from the GA-SVM algorithm was first evaluated on the EEG-locked on-scalp MEG IEDs, successfully classifying all on-scalp MEG IEDs that were not obscured by high-amplitude artifacts (cf. Fig. 4. Fig. 11). Running the GA-SVM on the remaining on-scalp MEG dataset resulted in the classification of 416 additional events as potential IEDs (Fig. 12). However, keeping only events constituting time series anomalies left only 31 events (Fig. 13–14). In comparison to in-helmet MEG, where MEG data showed 8 IEDs in addition to the 16 IEDs also seen by EEG, this demonstrates a potential increase in IED detection compared to EEG from 50% using in-helmet MEG to almost 200% using on-scalp MEG. Visual inspection of these 31 additional IEDs (Fig. 14) reveal a striking resemblance with the EEG-positive IEDs, showing that the algorithm-based IED detection discerns visually convincing IEDs. The resemblance between EEG-positive and algorithm-detected IEDs indicates a consistency in the statistical features underlying both categories of IEDs. Our results demonstrate a feasibility in registering and detecting IEDs in on-scalp MEG data, but also show that the added complexity in on-scalp MEG data might require assistance from algorithms so pick up on the abstract statistical features of IED events. In the in-helmet MEG data, the IEDs do not display this type of complexity and can readily be visually identified, why a GA-based approach is not needed or relevant for that data set.

### 4.4 Challenges and limitations

There are several limitations to this study. The ultimate value of on-scalp MEG epilepsy recordings can be said to depend on the extent to which on-scalp MEG can acquire information that is not available to in-helmet MEG or other existing non-invasive technologies. We present a successful benchmarking protocol that may be used to demonstrate the identification of IEDs uniquely detected by on-scalp MEG. However, the data consist of just one session from a single patient using a relatively small on-scalp MEG sensor array.

To further evaluate the potential usefulness of on-scalp MEG in epilepsy, as well as to evaluate the GA-SVM approach for IED detection, further studies are needed: preferably with larger-coverage (ideally whole-head) on-scalp MEG system, preferably on several epilepsy patients, and preferably with a higher-density co-registered EEG in both conventional and on-scalp MEG. The present study demonstrates that such studies are feasible, both from the perspective of screening suitable patients and from the perspective of a data recording protocol.

### 4.4 Conclusions

In this study, we present data from measurements on a temporal lobe epilepsy patient, where both on-scalp MEG data and in-helmet MEG data are obtained and compared. Using a benchmarking protocol aimed to quantify the amount of IEDs that are captured by on-scalp MEG, as compared to in-helmet MEG, we employed a novel automatic IED detection algorithm validated on the patient’s in-helmet MEG recording. The results indicate that we were able to find almost twice as many IEDs in the on-scalp MEG recording (42 IEDs: 16 EEG positive IEDs and 31 MEG-only IEDs) as we did in the in-helmet MEG measurement (24: 16 EEG positive IEDs and 8 MEG-only IEDs). It is possible that the additional IEDs detected in on-scalp MEG stem from cortical sources that are too small to be reflected in EEG or in-helmet MEG, potentially indicating that the on-scalp MEG system can identify IEDs that are not detectable by other non-invasive methods. Additional studies are needed to further evaluate the potential clinical usefulness of on-scalp MEG in epilepsy.

## References

Bartolomei, F., Lagarde, S., Wendling, F., McGonigal, A., Jirsa, V., Guye, M., Bénar, C., 2017. Defining epileptogenic networks: Contribution of SEEG and signal analysis. Epilepsia. https://doi.org/10.1111/epi.13791

Borna, A., Carter, T.R., Goldberg, J.D., Colombo, A.P., Jau, Y.Y., Berry, C., McKay, J., Stephen, J., Weisend, M., Schwindt, P.D.D., 2017. A 20-channel magnetoencephalography system based on optically pumped magnetometers. Phys. Med. Biol. 62, 8909–8923. https://doi.org/10.1088/1361-6560/aa93d1

Boto, E., Bowtell, R., Krüger, P., Fromhold, T.M., Morris, P.G., Meyer, S.S., Barnes, G.R., Brookes, M.J., 2016. On the potential of a new generation of magnetometers for MEG: A beamformer simulation study. PLoS One 11(8). https://doi.org/10.1371/journal.pone.0157655

Boto, E., Holmes, N., Leggett, J., Roberts, G., Shah, V., Meyer, S.S., Muñoz, L.D., Mullinger, K.J., Tierney, T.M., Bestmann, S., Barnes, G.R., Bowtell, R., Brookes, M.J., 2018. Moving magnetoencephalography towards real-world applications with a wearable system. Nature 555, 657–661. https://doi.org/10.1038/nature26147

Budker, D., Romalis, M., 2007. Optical magnetometry. Nat. Phys. 3, 227–234. https://doi.org/10.1017/CBO9780511846380

Chandola, V., Banerjee, A., Kumar, V., 2009. Anomaly detection: A survey. ACM Comput. Surv. 41, 1–58. https://doi.org/10.1145/1541880.1541882

Colon, A.J., Ossenblok, P., Nieuwenhuis, L., Stam, K.J., Boon, P., 2009. Use of routine MEG in the primary diagnostic process of epilepsy. J. Clin. Neurophysiol. 26, 326–332. https://doi.org/10.1097/WNP.0b013e3181baabef

Dale, A.M., Fischl, B., Sereno, M.I., 1999. Cortical Surface-Based Analysis Segmentation, I Reconstruction, Surface. Neuroimage 9, 179–194. https://doi.org/10.1006/nimg.1998.0395

De Tiège, X., Carrette, E., Legros, B., Vonck, K., Op De Beeck, M., Bourguignon, M., Massager, N., David, P., Van Roost, D., Meurs, A., Lapere, S., Deblaere, K., Goldman, S., Boon, P., Van Bogaert, P., 2012. Clinical added value of magnetic source imaging in the presurgical evaluation of refractory focal epilepsy. J. Neurol. Neurosurg. Psychiatry 83, 417–423. https://doi.org/10.1136/jnnp-2011-301166

De Tiège, X., Lundqvist, D., Beniczky, S., Seri, S., Paetau, R., 2017. Current clinical magnetoencephalography practice across Europe: Are we closer to use MEG as an established clinical tool? Seizure 50, 53–59. https://doi.org/10.1016/j.seizure.2017.06.002

Duez, L., Beniczky, S., Tankisi, H., Hansen, P.O., Sidenius, P., Sabers, A., Fuglsang-Frederiksen, A., 2016. Added diagnostic value of magnetoencephalography (MEG) in patients suspected for epilepsy, where previous, extensive EEG workup was unrevealing. Clin. Neurophysiol. 127, 3301–3305. https://doi.org/10.1016/j.clinph.2016.08.006

Fischl, B., Sereno, M.I., Dale, A.M., 1999. Cortical surface-based analysis: II. Inflation, flattening, and a surface-based coordinate system. Neuroimage 9, 195–207. https://doi.org/10.1006/nimg.1998.0396

Gonzalez-Martinez, J., Mullin, J., Bulacio, J., Gupta, A., Enatsu, R., Najm, I., Bingaman, W., Wyllie, E., Lachhwani, D., 2014. Stereoelectroencephalography in children and adolescents with difficult-to-localize refractory focal epilepsy. Neurosurgery. https://doi.org/10.1227/NEU.0000000000000453

Gramfort, A., Luessi, M., Larson, E., Engemann, D.A., Strohmeier, D., Brodbeck, C., Goj, R., Jas, M., Brooks, T., Parkkonen, L., Hämäläinen, M., 2013. MEG and EEG data analysis with MNE-Python. Front. Neurosci. 7, 1–13. https://doi.org/10.3389/fnins.2013.00267

Hari R, Baillet S, Barnes G, Burgess R, Forss N, Gross J, Hämäläinen M, Jensen O, Kakigi R, Mauguière F, Nakasato N, Puce A, Romani G-L, Schnitzler A, Taulu S. 2018. IFCN-endorsed practical guidelines for clinical magnetoencephalography (MEG). Clin. Neurophysiol. 129, 1720–1747. https://doi.org/10.1016/j.clinph.2018.03.042

Hämäläinen, M., Hari, R., Ilmoniemi, R.J., Knuutila, J., Lounasmaa Olli V., 1993. Magnetoencephalography - theory, instrumentation, and applications to noninvasive studies of the working human brain. Revies Mod. Phys. 65, 413–497.

Hämäläinen, M., Ilmoniemi, R.J., 1994. Interpreting magnetic fields of the brain: minimum norm estimates. Med. Biol. Eng. Comput. 32, 35–42.

Heiden, C., 1991. SQUID and SQUID system developments for biomagnetic applications. Clin. Phys. Physiol. Meas. 12, 67–73. https://doi.org/10.1088/0143-0815/12/B/009

Iivanainen, J., Stenroos, M., Parkkonen, L., 2017. Measuring MEG closer to the brain: Performance of on-scalp sensor arrays. Neuroimage 147, 542–553. https://doi.org/10.1016/j.neuroimage.2016.12.048

Iivanainen, J., Zetter, R., Grön, M., Hakkarainen, K., Parkkonen, L., 2019. On-scalp MEG system utilizing an actively shielded array of optically-pumped magnetometers. Neuroimage 194, 244–258. https://doi.org/10.1016/j.neuroimage.2019.03.022

Janca, R., Krsek, P., Jezdik, P., Cmejla, R., Tomasek, M., Komarek, V., Marusic, P., Jiruska, P., 2018. The sub-regional functional organization of neocortical irritative epileptic networks in pediatric epilepsy. Front. Neurol. 9, 1–11. https://doi.org/10.3389/fneur.2018.00184

Jayakar, P., Gaillard, W.D., Tripathi, M., Libenson, M.H., Mathern, G.W., Cross, J.H., 2014. Diagnostic test utilization in evaluation for resective epilepsy surgery in children. Epilepsia 55, 507–518. https://doi.org/10.1111/epi.12544

Jayakar, P., Gotman, J., Harvey, A.S., Palmini, A., Tassi, L., Schomer, D., Dubeau, F., Bartolomei, F., Yu, A., Kršek, P., Velis, D., Kahane, P., 2016. Diagnostic utility of invasive EEG for epilepsy surgery: Indications, modalities, and techniques. Epilepsia 57, 1735–1747. https://doi.org/10.1111/epi.13515

Jehi, L., 2018. The epileptogenic zone: Concept and definition. Epilepsy Curr. 18, 12–16. https://doi.org/10.5698/1535-7597.18.1.12

Jung, J., Bouet, R., Delpuech, C., Ryvlin, P., Isnard, J., Guenot, M., Bertrand, O., Hammers, A., Mauguière, F., 2013. The value of magnetoencephalography for seizure-onset zone localization in magnetic resonance imaging-negative partial epilepsy. Brain 136, 3176–3186. https://doi.org/10.1093/brain/awt213

Kane, N., Acharya, J., Benickzy, S., Caboclo, L., Finnigan, S., Kaplan, P.W., Shibasaki, H., Pressler, R., van Putten, M.J.A.M., 2017. A revised glossary of terms most commonly used by clinical electroencephalographers and updated proposal for the report format of the EEG findings. Revision 2017. Clin. Neurophysiol. Pract. 2, 170–185. https://doi.org/10.1016/j.cnp.2017.07.002

Keller, C.J., Truccolo, W., Gale, J.T., Eskandar, E., Thesen, T., Carlson, C., Devinsky, O., Kuzniecky, R., Doyle, W.K., Madsen, J.R., Schomer, D.L., Mehta, A.D., Brown, E.N., Hochberg, L.R., Ulbert, I., Halgren, E., Cash, S.S., 2010. Heterogeneous neuronal firing patterns during interictal epileptiform discharges in the human cortex. Brain 133, 1668–1681. https://doi.org/10.1093/brain/awq112

Knake, S., Halgren, E., Shiraishi, H., Hara, K., Hamer, H.M., Grant, P.E., Carr, V.A., Foxe, D., Camposano, S., Busa, E., Witzel, T., Hämäläinen, M.S., Ahlfors, S.P., Bromfield, E.B., Black, P.M., Bourgeois, B.F., Cole, A.J., Cosgrove, G.R., Dworetzky, B.A., Madsen, J.R., Larsson, P.G., Schomer, D.L., Thiele, E.A., Dale, A.M., Rosen, B.R., Stufflebeam, S.M., 2006. The value of multichannel MEG and EEG in the presurgical evaluation of 70 epilepsy patients. Epilepsy Res. 69, 80–86. https://doi.org/10.1016/j.eplepsyres.2006.01.001

Knowlton, R.C., Elgavish, R., Howell, J., Blount, J., Burneo, J.G., Faught, E., Kankirawatana, P., Riley, K., Morawetz, R., Worthington, J., Kuzniecky, R.I., 2006. Magnetic source imaging versus intracranial electroencephalogram in epilepsy surgery: A prospective study. Ann. Neurol. 59, 835–842. https://doi.org/10.1002/ana.20857

Luders, H.O., Iwasaki, M., Pataraia, E., H.O., L., M., I., E., P., 2004. Does magnetoencephalography add to scalp video-EEG as a diagnostic tool in epilepsy surgery? [4] (multiple letters). Neurology 63, 1987–1988.

Mitchell, M., 1998. An introduction to Genetic Algorithms, first ed., Cambridge, Massachusetts, Cambridge, Massachusetts, A Bradford Book The MIT Press

Murakami, H., Wang, Z.I., Marashly, A., Krishnan, B., Prayson, R.A., Kakisaka, Y., Mosher, J.C., Bulacio, J., Gonzalez-Martinez, J.A., Bingaman, W.E., Najm, I.M., Burgess, R.C., Alexopoulos, A. V., 2016. Correlating magnetoencephalography to stereo-electroencephalography in patients undergoing epilepsy surgery. Brain 139, 2935–2947. https://doi.org/10.1093/brain/aww215

Pataraia, E., Simos, P.G., Castillo, E.M., Billingsley, R.L., Sarkari, S., Wheless, J.W., Maggio, V., Maggio, W., Baumgarter, J.E., Swank, P.R, Breier, J.I., Papanicolaou, A.C., 2004. Does magnetoencephalography add to scalp video-EEG as a diagnostic tool in epilepsy surgery? [4] (multiple letters). Neurology 63, 1987–1988.

Pedregosa, F., Michel, V., Grisel, O., Blondel, M., Prettenhofer, P., Weiss, R., Vanderplas, J., Cournapeau, D., Pedregosa, F., Varoquaux, G., Gramfort, A., Thirion, B., Grisel, O., Dubourg, V., Passos, A., Brucher, M., Perrot, M., Duchesnay, A., 2011. Scikit-learn: Machine Learning in Python, Journal of Machine Learning Research.

Pfeiffer, C., Ruffieux, S., Jönsson, L., Chukharkin, M.L., Kalaboukhov, A., Xie, M., Winkler, D., Schneiderman, J.F., 2019. A 7-channel high-Tc SQUID-based on-scalp MEG system. bioRxiv 534107. https://doi.org/10.1101/534107

Rampp, S., Stefan, H., Wu, X., Kaltenhäuser, M., Maess, B., Schmitt, F.C., Wolters, C.H., Hamer, H., Kasper, B.S., Schwab, S., Doerfler, A., Blümcke, I., Rössler, K., Buchfelder, M., 2019. Magnetoencephalography for epileptic focus localization in a series of 1000 cases. Brain 0, 1–13. https://doi.org/10.1093/cercor/bhw393

Riaz, B., Pfeiffer, C., Schneiderman, J.F., 2017. Evaluation of realistic layouts for next generation on-scalp MEG: Spatial information density maps. Sci. Rep. 7. https://doi.org/10.1038/s41598-017-07046-6

Rosenow, F., Luders, H., 2001. Presurgical evaluation of epilepsy patients. Brain 124, 1683–1700. https://doi.org/10.1093/brain/124.9.1683

Sabolek, H.R., Swiercz, W.B., Lillis, K.P., Cash, S.S., Huberfeld, G., Zhao, G., Ste. Marie, L., Clemenceau, S., Barsh, G., Miles, R., Staley, K.J., 2012. A Candidate Mechanism Underlying the Variance of Interictal Spike Propagation. J. Neurosci. 32, 3009–3021. https://doi.org/10.1523/jneurosci.5853-11.2012

Schneiderman, J.F., 2014. Information content with low-vs. high-Tc SQUID arrays in MEG recordings: The case for high-Tc SQUID-based MEG. J. Neurosci. Methods. https://doi.org/10.1016/j.jneumeth.2013.10.007

Schoffelen, J.M., Gross, J., 2009. Source connectivity analysis with MEG and EEG. Hum. Brain Mapp. 30, 1857–1865. https://doi.org/10.1002/hbm.20745

Stefan, H., da Silva, F.H.L., 2013. Epileptic neuronal networks: Methods of identification and clinical relevance. Front. Neurol. 4 MAR, 1–15. https://doi.org/10.3389/fneur.2013.00008

Stefan, H., Hummel, C., Scheler, G., Genow, A., Druschky, K., Tilz, C., Kaltenhäuser, M., Hopfengärtner, R., Buchfelder, M., Romstöck, J., 2003. Magnetic brain source imaging of focal epileptic activity: A synopsis of 455 cases. Brain 126, 2396–2405. https://doi.org/10.1093/brain/awg239

Sutherling, W.W., Mamelak, A.N., Thyerlei, D., Maleeva, T., Minazad, Y., Philpott, L., Lopez, N., 2008. Influence of magnetic source imaging for planning intracranial EEG in epilepsy. Neurology 71, 990–996. https://doi.org/10.1212/01.wnl.0000326591.29858.1a

Tao, J.X., Ray, A., Hawes-Ebersole, S., Ebersole, J.S., 2005. Intracranial EEG Substrates of Scalp EEG Interictal Spikes. Epilepsia 45, 669–676.

Taulu, S., Simola, J., 2006. Spatiotemporal signal space separation method for rejecting nearby interference in MEG measurements. Phys. Med. Biol. 51, 1759–1768. https://doi.org/10.1088/0031-9155/51/7/008

Wilke, C., Worrell, G.A., He, B., 2011. Graph analysis of epileptogenic networks in human partial epilepsy. Epilepsia 52, 84–93. https://doi.org/10.1038/mp.2011.182.doi

Wilson, S.B., Emerson, R., 2002. Spike detection: A review and comparison of algorithms. Clin. Neurophysiol. 113, 1873–1881. https://doi.org/10.1016/S1388-2457(02)00297-3

Xie, M., Schneiderman, J.F., Chukharkin, M.L., Kalabukhov, A., Riaz, B., Lundqvist, D., Whitmarsh, S., Hämäläinen, M., Jousmäki, V., Oostenveld, R., Winkler, D., 2017. Benchmarking for on-scalp MEG sensors. IEEE Trans. Biomed. Eng. 64, 1270–1276. https://doi.org/10.1109/TBME.2016.2599177

Xie, M., Schneiderman, J.F., Chukharkin, M.L., Kalabukhov, A., Whitmarsh, S., Lundqvist, D., Winkler, D., 2015. High-Tc SQUID vs. low-Tc SQUID-based recordings on a head phantom: Benchmarking for magnetoencephalography. IEEE Trans. Appl. Supercond. 25. https://doi.org/10.1109/TASC.2014.2366420

Zhang, L., Liang, Y., Li, F., Sun, H., Peng, W., Du, P., Si, Y., Song, L., Yu, L., Xu, P., 2017. Time-Varying Networks of Inter-Ictal Discharging Reveal Epileptogenic Zone. Front. Comput. Neurosci. 11, 1–9. https://doi.org/10.3389/fncom.2017.00077

Zhang, Y., Tavrin, Y., Mück, M., Braginski, A.I., Heiden, C., Hampson, S., Pantev, C., Elbert, T., 1993. Magnetoencephalography using high temperature rf SQUIDs. Brain Topogr. 5, 379–382. https://doi.org/10.1007/BF01128694

