## Supplementary for "Detection of interictal epileptiform discharges: A comparison of on-scalp MEG and conventional MEG measurements"

### 6 Supplementary

*6.1 Combined genetic algorithm and support vector machine*

A genetic algorithm (GA) aims to solve an optimization problem by testing a group ("population") of candidate solutions ("chromosomes") using a fitness function to evaluate the individual solutions. Good solutions ("parents") are kept and allowed to exchange information ("crossover") producing a new generation ("children"). Random changes ("mutations") are introduced in these, producing new chromosomes to be tested by the fitness function. A large variety of solutions can be tested until the most optimal one is found (Mitchell, 1998). Here, chromosomes were feature vectors tested by a fitness function determining the overall similarity between the chromosome and the EEG-locked on-scalp MEG IEDs.

6.1.1 Creation of the first generation of candidate solutions

Initially, on-scalp MEG raw data epochs (0.4 seconds) coinciding with IEDs in the EEG were constructed and averaged across sensors. Suppose $n$ such epochs were found. The 14 feature $f_i, 1 \leq i \leq 14$, of Table 2 were extracted from these, resulting in $n$ *IED feature vectors*, $[f_1^{(j)}, ..., f_{14}^{(j)}], 1 \leq j \leq n$. The initial candidate solutions (the first generation of chromosomes), $c^{(k)}, 1 \leq k \leq 10000$, 10000 being the size of this population, were defined as $c^{(k)} = [g_1^{(k)}, ..., g_{14}^{(k)}]$, with $g_i^{(k)}, 1 \leq i \leq 14$, randomly drawn from the interval $[\min f_i^{(j)}, \max f_i^{(j)}]$.

6.1.2 Features

The following 14 features were extracted from the on-scalp MEG raw data epochs:

| |
|---|
| **Standard deviation** |
| **Skewness** |
| **Mean** |
| **Kurtosis** |
| **Sum of time points in epoch** , $\Sigma_{i=1}^{N} x_i$, $x_i$ the $i$:th data point, $N$ the number of data points in the epoch. |
| **Maximum value of epoch** |
| **Minimum value of epoch** |
| **Range of epoch** |
| **Energy of epoch**, $\frac{1}{N}\Sigma_{i=1}^{N} x_i^2$ |
| **Integral of epoch** |
| **Duration of peak** |
| **Fractal dimension** $D$, $D = \frac{\log_{10}(L)}{\log_{10}(d)}$, $L$ the total length of the curve calculated as the Euclidean distance between samples; $d$ the maximum distance between the first data point and any other point. |
| **Variance** |
| **Slope of peak** |

Table 2 Features extracted from EEG-positive on-scalp MEG IEDs used to create artificial IED feature vectors

6.1.3 Fitness function

The fitness function tested the similarity, $s_i^{(k)}$, between the $i$:th feature of the $k$:th chromosome and the corresponding feature of the IED feature vectors according to

$$s_i^{(k)} = \sqrt{(g_i^{(k)} - f_i^{(1)})^2 + (g_i^{(k)} - f_i^{(2)})^2 + ... + (g_i^{(k)} - f_i^{(n)})^2}$$

For each chromosome the vector $[s_1^k, ..., s_{14}^k]$ was acquired. The fitness value $ft^{(k)}$, of the $k$:th chromosome $c^{(k)}$ was defined as the $\ell_2$-norm of this vector,

$$ft^{(k)} = \|[s_1^k, ..., s_{14}^k]\|$$

Thus, the smaller $ft^{(k)}$, the greater the resemblance between chromosome $c^{(k)}$ and all of the IED feature vectors.

6.1.4 Crossover and mutation

Parents were chosen using binary tournament selection (for details see (Mitchell, 1998)). Crossover was performed by exchange of features between parents, and crossover rate was set to 0.5. Resulting children mutated with mutation rate $\frac{1}{\text{chromosome length}}$. The resulting mutated feature, $g_i^{(k)}(mut)$, was defined as $g_i^{(k)}(mut) = g_i^{(k)} + mut$ where $mut \sim \mathcal{N}(\mu, \sigma)$, $\mu = g_i^{(k)}$, $\sigma = \frac{|g_i^{(k)}|}{1300}$. In order to keep the best chromosomes, the 2000 parents with the best fitness value were passed on to the next generation without mutation. In all, 100 generations were created and the chromosome with the lowest fitness value were kept. 1000 such chromosomes were collected, labeled *artificial IED feature vectors* and used as training data for the support vector machine (SVM).

6.1.5 Creation of non-IED feature vectors

*Non-IED feature vectors* to also be used as training data were derived from a portion of the raw data without EEG-positive on-scalp MEG IEDs. The data was averaged across sensors and subdivided into 0.4 seconds long intervals. For each of these, the time point of maximum amplitude was located and new epochs centered around this point were created in order to capture non-IED waveforms. These epochs were visually inspected in order to exclude events resembling IEDs. The 14 features $f_i, 1 \leq i \leq 14$, were extracted from these epochs, forming non-IED feature vectors and 1000 such vectors were used. Python module Scikit-learn SVM with a RBF kernel (Pedregosa et al., 2011) was trained upon the artificial IED feature vector and the non-IED feature vectors. The trained SVM was validated on the EEG-locked on-scalp MEG IEDs, and classification was performed on remaining data set not used in training. After the initial classification based upon all 14 features, Scikit-learn feature selection SelectKBest using ANOVA F-value (Pedregosa et al., 2011) was performed in order to determine the $m$ most influential features. The seven features of Table 3 performed the best when validated on the EEG-positive IEDs. The GA-SVM was re-run using only these features. The events classified as IEDs using the seven features of Table 3 were labeled *potential IEDs* .

| |
|---|
| **Standard deviation** |
| **Skewness** |
| **Mean** |
| **Kurtosis** |
| **Sum of time points in epoch** |
| **Minimum value of the epoch** |
| **Integral of epoch** |

Table 3 Best performing features after validation on EEG-positive on-scalp MEG IEDs

*6.2 Anomaly detection*

Since an IED, by definition, constitutes a time series anomaly (Chandola et al., 2009; Kane et al., 2017), only non-stationary potential IEDs should be kept. To single out such events, a baseline epoch starting at 1.15 seconds before and ending at 0.2 seconds before maximum positive peak of the event and an event epoch starting at 0.75 seconds before and ending 0.2 seconds after the maximum positive peak of the event were created for the EEG-locked on-scalp MEG IEDs as well as for the potential IED-events. The seven best performing features (Table 3) were extracted from EEG-positive on-scalp MEG IED event and baseline epochs, $f_i^{(j)}$(baseline) and $f_i^{(j)}$(event), $2 \leq i \leq 7$, $1 \leq j \leq n, n$ = number of EEG-positive on-scalp MEG IED epochs, and from potential IEDs, $g_i^{(l)}$(baseline) and $g_i^{(l)}$(event), $2 \leq i \leq 7$, $1 \leq l \leq r, r =$ number of potential IEDs. For each feature, $f_i$, $f_i^{(j)}$(diff) was defined as the Euclidean distance between $f_i^{(j)}$(baseline) and $f_i^{(j)}$(event). For each feature, $f_i$, $f_i(\text{min}) = \min(f_i^{(1)}(\text{diff}), f_i^{(2)}(\text{diff}), ..., f_i^{(n)}(\text{diff}))$ was determined. Similarly, $g_i^{(l)}(\text{diff}), 1 \leq l \leq r$, was calculated. Potential IEDs fulfilling $g_i^{(l)}(\text{diff}) \geq f_i(\text{min})$ for all $i$ were kept and labeled *likely IEDs.*
